# Alpha cell inflammation during human pancreas aging and type 2 diabetes and its reversal by calorie restriction in mice

**DOI:** 10.1101/2025.06.04.657914

**Authors:** Michael Schleh, Jean-Philippe Cartailler, Shristi Shrestha, Amanda Cambraia, Aliyah Habashy, Cara Ellis, Melanie Cutler, Gabriel Ferguson, Alvin Powers, Rafael Arrojo e Drigo

## Abstract

Aging increases risk for type 2 diabetes (T2D), partly through loss of beta cell identity and function driven by metabolic stress and islet inflammation. While calorie restriction (CR) promotes beta cell longevity in young animals, its impact on cellular aging and inflammatory burden in older individuals is unclear. Using SCENIC regulon and multiomic analyses, we find that aging human alpha cells adopt a coordinated inflammatory phenotype marked by IFN-γ signaling to increase major histocompatibility complex (MHC) class I presentation, and CD8^+^ T cell recruitment and activation towards islets. In T2D, CD8^+^ T cells further progress towards an effector memory state. CR reduces alpha cell MHC-I expression while subsequently suppressing CD8^+^ effector status and accompanied by reduced islet inflammation and immune cell infiltration in mice. Together, these findings highlight an alpha cell–immune signaling axis in aging and T2D that may underlie fibrosis and disease pathophysiology.

## Introduction

Aging is a major risk factor for type 2 diabetes (T2D), with prevalence reaching 28.8% among adults aged ≥65 years in the United States (*1*). Age-related deterioration in glycemic control has largely been attributed to progressive insulin resistance, impaired beta cell insulin secretion, and altered hepatic insulin clearance (*2–4*). However, aging is also characterized by tissue loss of homeostatic resilience and increased immune heterogeneity, that may shape metabolic health decline (*5*). A central feature of adaptive immune aging is thymic involution, which reduces naïve T cell output and shifts the T cell compartment towards peripheral expansion and memory biased populations (*6–8*). Whether such age-associated immune remodeling arises primarily through intrinsic programs or occurs secondary to extrinsic cues in the microenvironment and whether these aspects are present during diabetes, remains largely unknown.

Pancreatic islets are composed primarily of long-lived endocrine alpha- and beta cells, which maintain glucose homeostasis through coordinated secretion of glucagon and insulin (*9–12*). During aging, beta cells exhibit structural and functional decline, including loss of cellular identity, DNA damage accumulation, endoplasmic reticulum (ER) stress, impaired autophagy, increased transcriptional noise, amyloid deposition, senescence, and accumulate islet fibrosis (*2, 13–16*). In contrast, far less is known how aging affects alpha cell biology despite well-known disruptions in alpha cell glucagon secretion during aging and along the natural history of T2D (*2, 17–20*) that are accompanied by exaggerated amino acid-induced glucagon release (*21, 22*). Moreover, it remains unknown whether aging promotes intrinsic inflammatory programs within specific endocrine cell types, or whether islet inflammation is driven predominantly by extrinsic immune cues. Defining the cellular origins of age-associated islet inflammation is essential to understanding the intersection of immune and metabolic aging and how these aspects may contribute to diabetes pathogenesis.

Calorie restriction (CR) is a well-established dietary intervention that extends lifespan and health span across species by delaying several hallmarks of cellular dysfunction during aging (*23–26*). Moderate energy restriction (10–40%) improves metabolic homeostasis in both young and aged animals, in part through modulation of nutrient-sensing pathways including insulin-IGF1, mTORC1, and AMPK (*27, 28*). CR also reduces oxidative stress and inflammatory tone while enhancing autophagy and proteostasis (*29–31*), and it can promote anti-inflammatory responses in lean adult humans (*32*). In young adult mice, CR and CR-mimicking interventions improve beta cell function and longevity, and which are thought to be explained by enhanced peripheral insulin sensitivity (*33–36*). However, whether CR can reverse established age-associated islet inflammation, particularly within specific endocrine cell populations, remains unresolved.

Here, we integrated multimodal and multiomic analyses of publicly available human islet datasets – including bulk proteomics, singe-cell RNA sequencing (scRNA-seq) and epigenome profiling, and spatial proteomics – and performed a large-scale meta-analysis to define the molecular signatures of alpha cells during aging in non-diabetic (ND) donors and to determine their relevance to T2D. At the islet proteome level, human islets exhibited hallmarks of ER stress, senescence, and inflammatory activation linked to innate and adaptive signaling compartments. Single cell transcriptomics revealed these inflammatory signatures were significantly enriched in alpha cells and these resulted from IFN-γ-induced pro-inflammatory signaling via activation of pro-inflammatory transcription factors (TFs) STAT1, IRF1, and NFKB1. Notably, this pro-inflammatory alpha cell state is linked to increased amino-acid induced glucagon release in vitro, which may explain previous observations of hyperglucagonemia in ND humans in vivo (*2*). Moreover, spatial proteomics confirmed this pro-inflammatory signaling in alpha cells correlates with increased abundance of major histocompatibility complex class I (MHC-I; HLA-ABC) in alpha cells from aged donors, and which associated with enrichment of CD8^+^ T cells within the islet microenvironment and closer to alpha cells, specifically. In T2D, these islet CD8+ T cells suffer a shift towards central memory and terminally differentiated CD8^+^ phenotypes. Similarly, we show in old mice that this alpha cell-immune association is active, and it can be successfully reversed by short term CR, which is sufficient to reduce alpha cell inflammation and attenuate CD8⁺ T cell effector islet accumulation.

Together, our study identifies alpha cell inflammation as a previously unrecognized hallmark of human pancreas aging that is molecularly and functionally distinct from aging beta cells. These findings establish a link between alpha cell pro-inflammatory programs and adaptive immune activation during aging and demonstrate that CR can reverse age-associated islet immune dysfunction by modulating alpha cells and to preserve systemic metabolic homeostasis.

## Results

### Aging ND islets exhibit elevated immune activity and unfolded protein response signaling

Islet inflammation and fibrosis are hallmarks of human pancreas aging and diabetes, contributing to beta cell dysfunction (*37–39*). To define proteomic signatures of aging, we leveraged the HumanIslets consortium dataset comparing young (28±4 years, n=26) and old (71±3 years, n=19) ND donors (**Figure 1A, Supplementary Table S1A-S1B**) (*40, 41*). Aging was associated with 86 increased and 17 decreased proteins (**Supplementary Table S2**). Pathway analysis (Reactome) revealed enrichment of innate immune signaling, alongside elevated amino acid metabolism, mTORC1 activity, and ER stress pathways in old adults (**Figure 1B, Supplementary Table S3**). Consistent with these changes, aged islets exhibited increased markers of beta cell immaturity (RBP4 (*42*); **Figure 1C**), senescence (CDKN1A/p21 (*43*), **Figure 1D**), ER stress signaling (ATF6, CHOP, PERK; **Figure 1E**), and mTORC1 activation (LAMP2, LAMTOR, RAGA, RAGC; **Figure 1F**). A robust immune signature was also evident by upregulation of antigen presentation (HLA-A), T-cell receptor (TCR) signaling and memory proteins (CD28, CD44), and immune-regulatory signaling (TGF-β1, TOM1, CD63; **Figure 1G**). Together, these findings reinforce aging human islets exhibit coordinated ER stress and metabolic remodeling alongside heightened inflammatory and immune engagement within the islet microenvironment.

**Figure 1.**
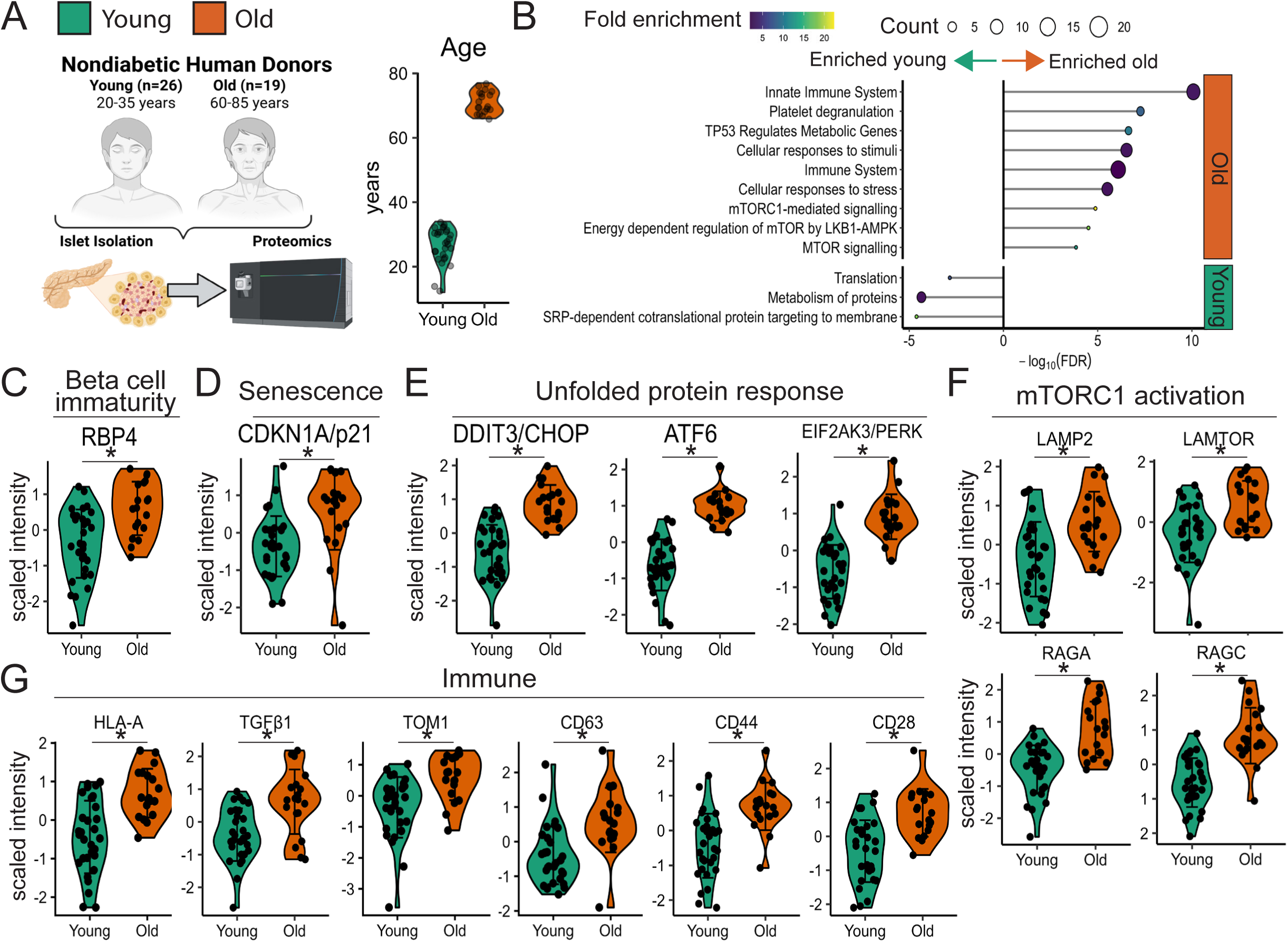
Aging human islets have an inflammatory activation phenotype combined with mTORC1 signaling and elevated unfolded protein response stress. **(A)** Islets from young (28±4 years, n=26), and old (71±3 years, n=19) ND donors from the HumanIslets consortium were compared by proteomics meta-analysis. **(B)** Reactome pathways present from significantly abundant proteins between young and old ND donors. **(C-G)** Protein expression levels for beta cell immaturity marker, RBP4 (C); senescence, CDKN1A/p21 (D); unfolded protein response, CHOP (DDIT3), ATF6, PERK (EIF2AK3) (E); mTORC1 activation, LAMP2, LAMTOR, RAGA, RAGC (F); and immune regulatory components, HLA-A, TGF-β1, TOM1, CD63, CD44, and CD28, respectively (G). HLA, human leukocyte antigen. *p<0.05, young vs. old donors. Error bars are mean ± SD.

### Single cell transcriptomics reveal inflammatory signaling in aging human alpha cells

Over the past decade, single-cell approaches have defined how islet gene regulation changes during development, aging, and diabetes (*42, 44–53*). We previously showed that aging ND beta cells exhibit chronic ER stress and loss of identity and insulin secretion (*44*), and further supported by our proteomic findings (**Figure 1**). To identify specific cell types contributing to the inflammatory signatures observed in aging islets, we analyzed scRNA-seq data from a uniformly re-processed dataset created by PanKbase based on previously published studies (*54, 55*). After quality control and cohort selection, the final dataset annotated 131,610 islet cells from 32 donors (56,851 alpha cells), stratified into young (20-40 years) and old (≥50 years) groups **(Supplementary Figure S1A-C, Supplementary Table S4A-S4B).**

Among ND islet endocrine cell types, alpha cells exhibited the highest expression of MHC-I genes (HLA-ABC), followed by immune, ductal, and endothelial cells (**Supplementary Figure S1D**). Pseudobulk differential expression and gene set enrichment analysis (GSEA) across alpha, beta, acinar, and MUC5B ductal cells identified age-dependent transcriptomic changes for major islet and pancreas cell types (**Figure 2B, Supplementary Figure S1E, Supplementary Table S5**). This approach revealed activation of type II interferon (IFN-γ) signaling in aging alpha, beta, and acinar cells (**Figure 2C and Supplementary Figure S1F**). To deconvolve the impact of IFN-γ within the human islet cytoarchitecture, we performed weighted analysis of IFN-γ responsive genes enriched in old alpha cells and beta cells and discovered that canonical IFN-γ signaling is strongly enriched in old alpha cells (**Figure 2D, Supplementary Figure S2A-S2B**). Old alpha cells displayed canonical IFNγ signatures, including activation of JAK-STAT (*STAT1*, *SOCS1*, *SOCS3*, *IRF7/9*, *IL7*, and *IL15*) and antigen processing pathways (*TAP1, B2M*), whereas old beta cells display TNF-signaling pathway signatures (**Figure 2D-E, Supplementary Figure S2C-S2D**). As expected, this analysis also found activation of ER stress and DNA damage, and impaired oxidative phosphorylation in old beta cells (*44*). None of these are found in old alpha cells (**Supplementary Figure S1F, S2A-S2B, Supplementary Table 6**). Old alpha cells also increased expression of senescence-associated genes (*IGF1R, CDKN2A, CDKN1A/B*) and of alpha cell identity markers (*ALDH1A1, TM4SF4, LOXL4)*, while displaying reduced expression of ER stress-related genes (HERPUD1, ATF4, JUND). This data hints that aging alpha cells may develop senescence next to old beta cells however they do not suffer from significant impairments to identity or activation of ER stress (**Supplementary Figure S1G**) (*56, 57*). Importantly, unlike old acinar cells, old alpha cells and beta cells did not display significant activation of pro-apoptosis pathways (**Supplementary Figure S1H**), thus suggesting that this pro-inflammatory environment does not lead to loss of alpha or beta cells.

**Figure 2.**
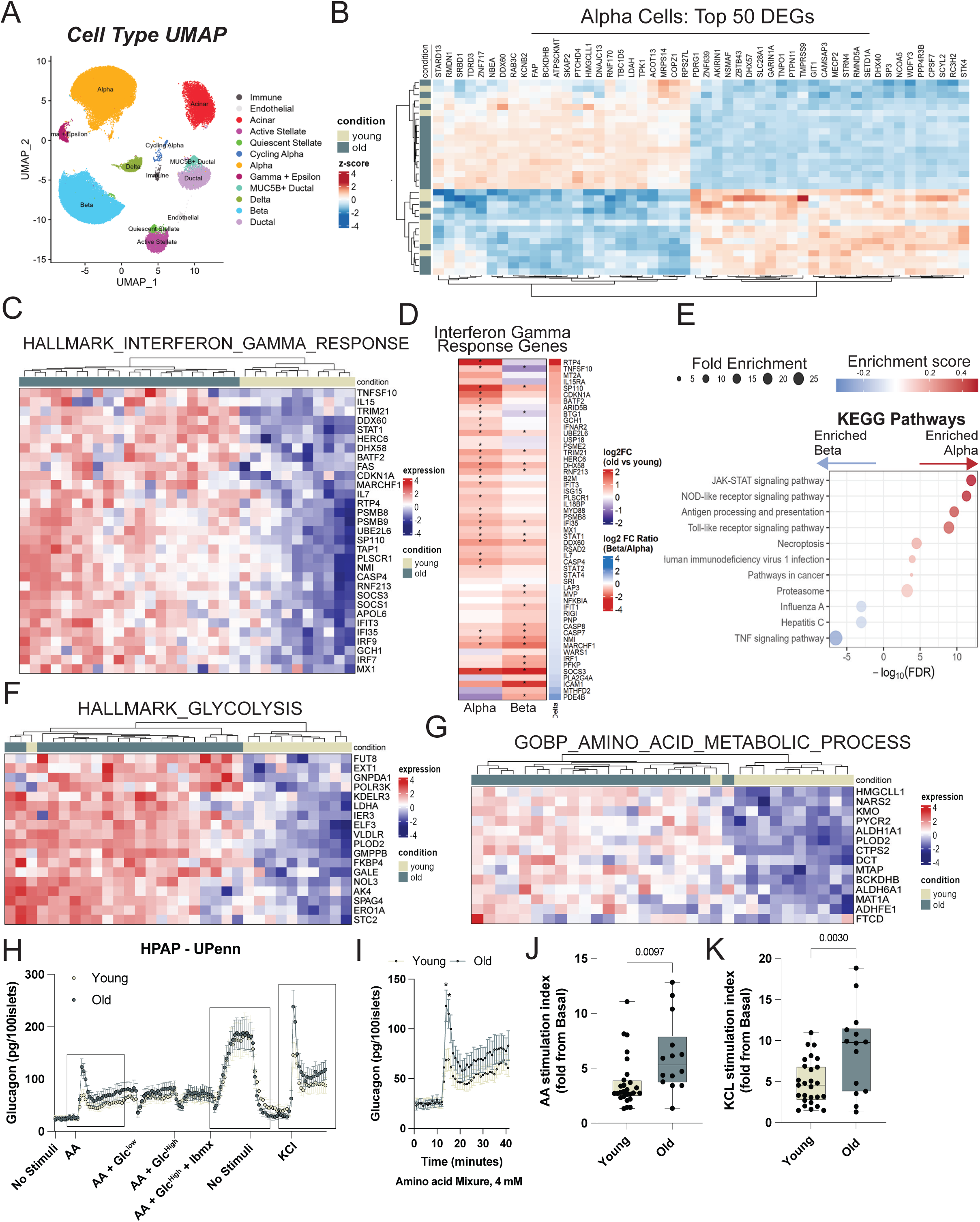
Aging increases human alpha cell interferon signaling and metabolic signatures. **(A)** Cell type UMAP of integrated scRNA-seq data obtained from HPAP database. **(B)** Heatmap with hierarchical clustering showing top 25 genes differentially regulated by age in human alpha cells. **(C)** GSEA hallmarks for gene sets associated with the Hallmark Interferon GAMMA Response (M5913). **(D)** Relative gene expression levels of genes in the Hallmark Interferon GAMMA Response (M5913) collection in alpha cells versus beta cells. Asterisks mark genes significantly upregulated in either cell type. **(E)** Weighted KEGG pathway analysis demonstrating the enrichment of IFN-γ associated genes in old alpha cells versus old beta cells. **(F-G)** Significantly expressed Hallmark genes for metabolic processes including Glycolysis (M5937; G) and Amino Acid Metabolic Process (GO:0006520), respectively. **(H-K)** Isolated human islet perifusion assay including stimulation with amino-acids, glucose, IBMX, and KCL. Dotted line boxes and represented quantification (**I**, **J**, and **K**) highlight specific stimuli that provoked significant glucagon secretion from old islets versus young islets. P-values are shown. Data are mean ± SD with boxplots representing mean ± IQR. ND, non-diabetic. HPAP, Human Pancreas Analysis Program; GSEA, gene set enrichment analysis.

Aging alpha cells also had increased expression of genes across several metabolic processes including glucose metabolism and oxidative phosphorylation, fatty acid metabolism, and amino acid catabolism (**Figure 2F-G and Supplementary Figure S1I**). To determine if these transcriptional changes correlate with alterations in alpha cell glucagon secretion, we analyzed HPAP human islet perifusion data from young and old adult donors from two different human islet analysis centers (UPenn and Vanderbilt), which used complementary stimulation protocols to test for the effect of several insulin and glucagon secretagogues. This revealed that alpha cells from older donors have increased glucagon secretion in response to stimulation with a mixed amino acid (AA) cocktail and to de-polarization with KCL (**Figure 2H-K**). No changes were observed when alpha cells were stimulated with low glucose, high glucose, and epinephrine (**Supplementary Figure S3**).

Together, these results indicate that aging human alpha cells acquire senescence-associated features, display heightened AA-dependent glucagon secretion, and an active IFN-γ signaling coupled with MHC-I antigen-presentation programs.

### Re-organization of gene regulatory networks activates inflammatory gene expression in aging alpha cells

We have previously applied the transcription factor (TF) inference algorithm pySCENIC (*58*) to show that aging beta cell stress arises from reconfigured gene regulatory networks (GRNs) and activation of stress TFs (*44*). Here, we used pySCENIC and graph network analysis to characterize GRNs in aging alpha cells and identify the TFs linked to activation of the observed inflammatory response. This approach identified TFs enriched in both age groups (**Figure 3A**), including canonical alpha cell TFs (NKX6-1,FOXA2), and inflammation-associated TFs that were significantly enriched in old alpha cells (IRFs, NFKB1, and STAT1) (**Figure 3A, Supplementary Figure S4**).

**Figure 3.**
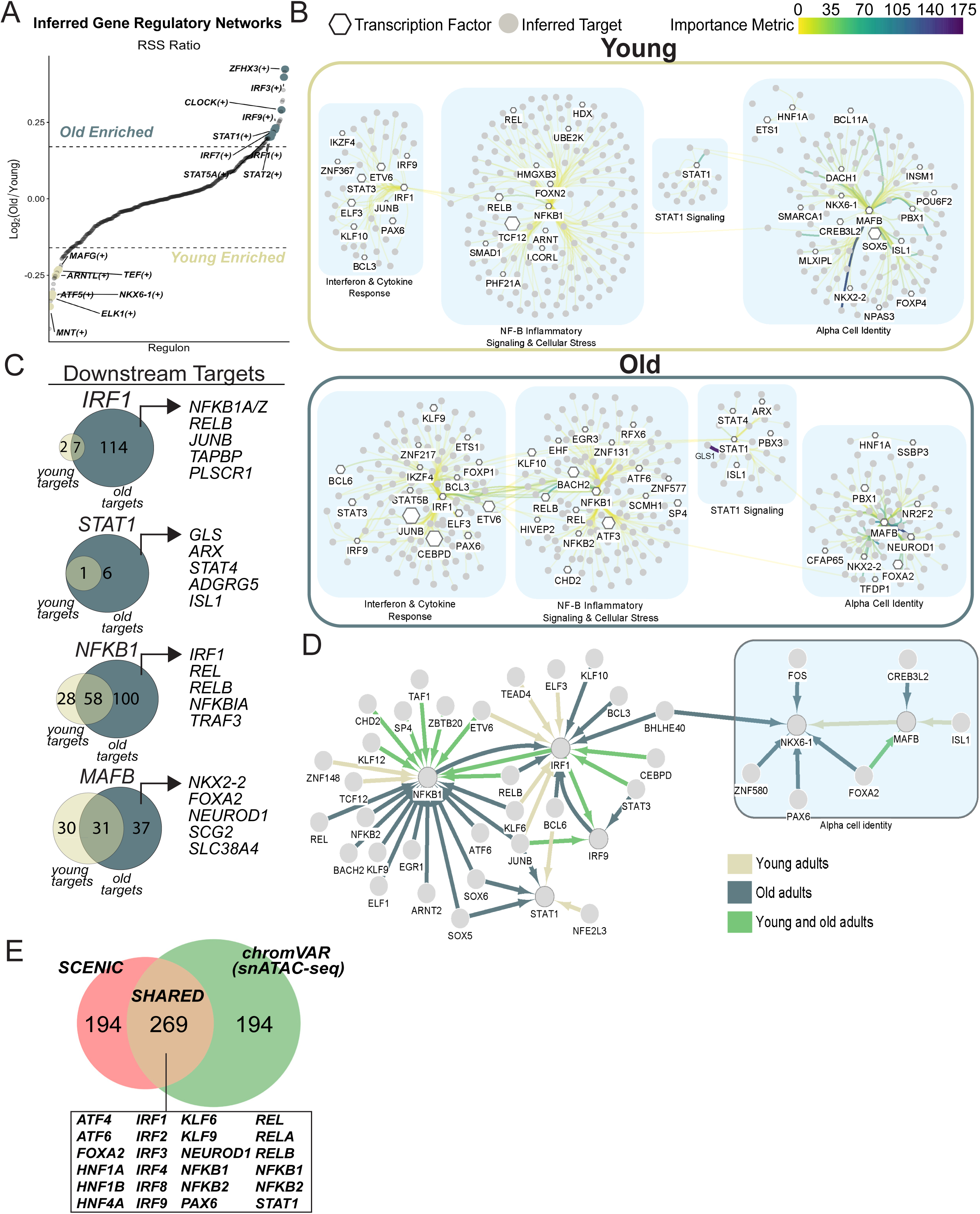
Re-wiring of gene regulatory networks activates an interferon-stimulated inflammatory phenotype in aging alpha cells. **(A)** Regulon specificity scores (RSS) ratios between old and young alpha cells, with enrichment defined as the top 10% of RSS ratios within each group. **(B)** Annotated Gene Regulatory Network (GRN) maps of young (top) and old (bottom) human alpha cells. Transcription factors (TFs) are represented by white hexagons and inferred TF target genes are shown in solid grey circles. Only the top 15% TF regulons after 100X pySCENIC inference simulations are shown. TF hexagon size represents the TF importance metric within the GRN structure, and TF-target gene nodes are connected by line edges colored using a *viridis* hue to represent predicted connection strength. Shaded blue boxes are annotated with pathway enrichment terms associated with their respective regulon network. **(C)** Venn diagrams representing unique and shared genes found in the regulons associated with IRF1, NFKB1, STAT1, AND MAFB TFs. **(D)** Regulatory network map displaying age-dependent and age-independent regulatory connections between inflammatory TFs and canonical alpha cell identity TFs. In this map, TFs are represented by solid grey circles and regulatory relationships are shown by arrows. Green arrows, connections found in both young and old alpha cells; dark blue and beige arrows represent connections found uniquely in old and young alpha cells, respectively. **(E)** Venn diagram illustrating the overlap between SCENIC and chromVAR associated TFs detected in old alpha cells.

We reconstructed enriched GRNs of young and old alpha cells and observed a stronger inferred activity of pro-inflammatory TFs (IRF1, NFKB1, and STAT1 signaling) and reduced activity of MAFB during aging (**Figure 3B**). This data suggests that although inflammation-associated TFs are found in both age groups, these TFs play a relatively small role in the overall transcriptome landscape in younger alpha cells. To test this hypothesis, we performed analysis of TF-associated regulons to reveal that inflammatory TFs do not target the same set of genes in young versus old alpha cells, suggesting a significant re-wiring of alpha cell GRNs during aging. This is characterized by a low Jaccard similarity index between conserved TFs in both age groups, and this includes loss of TF-regulon similarity for alpha cell identity TFs (ARX, HNF1A, HNF4A) as well as inflammation-associated TFs (STAT1, IRF2/7/8/9). In fact, these TFs are redirected towards new targets that include several genes identified as part of the pro-inflammatory signaling phenotype of old alpha cells (**Figure 3C**, **Supplementary Table S7**). Moreover, several of these downstream gene targets were pro-inflammatory TFs themselves, which created a pro-inflammatory TF network in old alpha cells centered around NFKB1 and IRF1 (**Figure 3D**). In contrast, old alpha cells lose the link between NKX6.1, MAFB, and ISL1 that is only observed in young cells (**Figure 3D**).

To further validate our SCENIC results, we integrated our analysis with publicly available chromatin accessibility data from HPAP (snATAC-seq). Here, we performed differential analysis of peak accessibility using ChromVAR (*59*) to identify TF motifs enriched in chromatin regions significantly upregulated in old alpha cells. Next, we crossed referenced this list with our old alpha cell SCENIC data to identify n=269 shared TFs, including canonical alpha cell TFs (FOXA2, HNF1A/B) and several inflammatory TFs (IRFs, KLF6, NFKB1/RELs, STAT1) (**Figure 3E**).

These data indicate that aging promotes re-wiring of alpha cell GRNs to activate interferon-stimulated NFKB1, IRF1, and STAT1 gene expression programs and this likely explains the origins of pro-inflammatory activation in aging alpha cells.

### Spatial analysis of aging human pancreases identifies alpha cell-T-cell interactions *in situ*

Aging ND human pancreases exhibit multiple features of chronic low-grade inflammation, including increased fibrosis, higher incidence of acinar-to-ductal metaplasia (ADM), angiopathy, and pancreatic adiposity (*38, 39*). To validate our proteomic and transcriptomics meta-analysis findings *in situ*, we analyzed high-dimensional imaging mass cytometry (IMC) and CO-Detection by indexing (CODEX) datasets from HPAP (*60, 61*). Human ND pancreas samples were stratified into younger adult (22-40 years) and older adults (47-58 years) groups (**Supplementary Figure S5B and S6B; Supplementary Tables S8 and S9).** Individual cells from each donor image were segmented using standard QuPath- and Seurat-based workflows for cell identification and spatial annotation, cell marker quantification, and unsupervised clustering (**Supplementary Figure S5A and S6A)**. This allowed us to identify and map the spatial landscape of major pancreatic cells including alpha cells (GCG^+^), beta cells (C-peptide^+^), delta cells (SST^+^), pancreatic polypeptide cells (PP^+^), endothelial cells (CD31^+^), Epithelial/Acinar cells (CK19^+^/CA2^+^), M1-like macrophages (CD45^+^CD68^+^CD163^-^), M2-like macrophages (CD45^+^CD68^+^CD163^+^), and CD4^+^ and CD8^+^ T cell subpopulations (CD45^+^CD3^+^CD4^+^ and CD45^+^CD3^+^CD8^+^, respectively) (**Figure 4A and 4C, Supplementary Figures S5C-G and S6C-G, Supplementary Tables S10-S11**).

**Figure 4.**
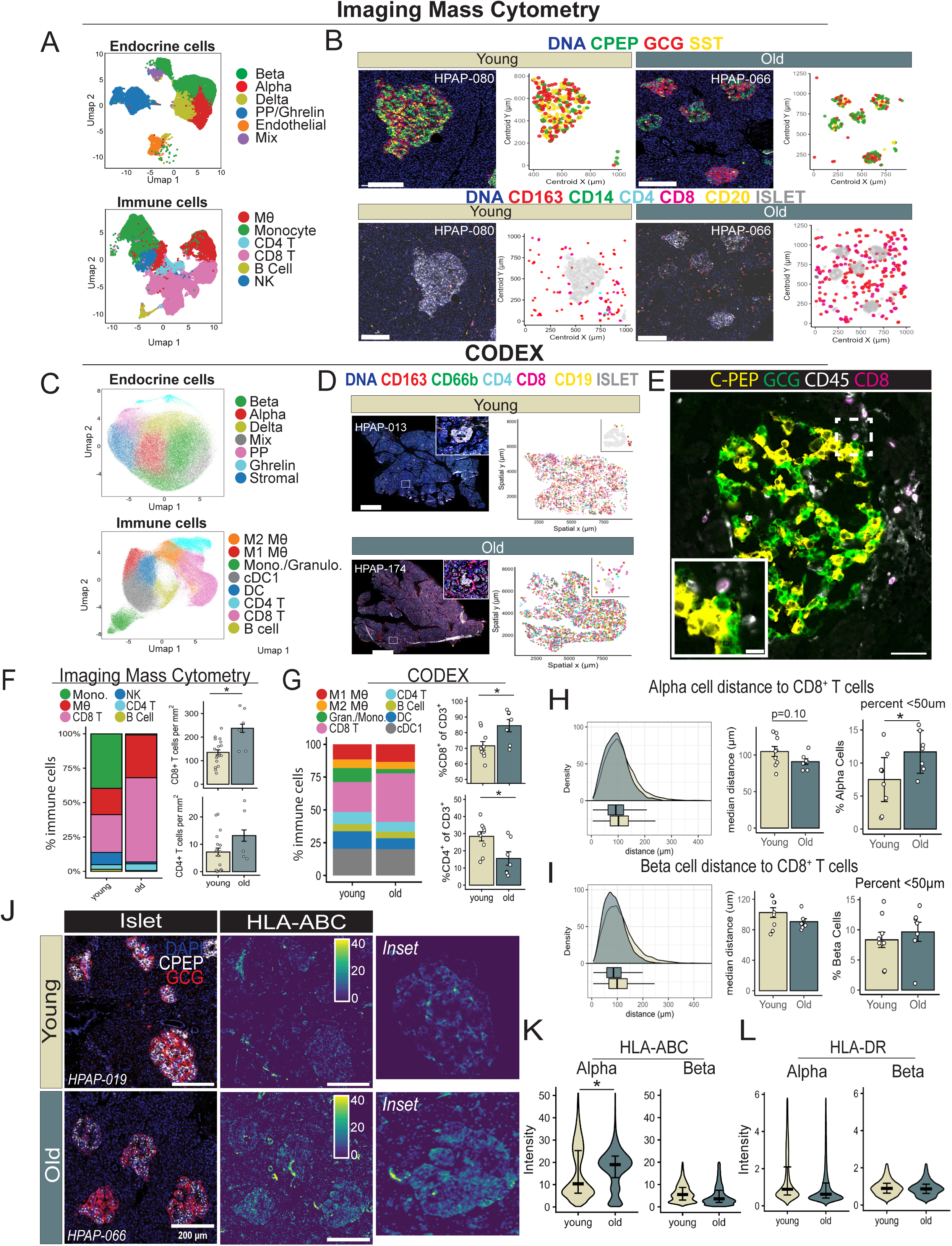
Aging increases CD8^+^ islet infiltration via alpha cell MHC-I signaling. **(A)** Imaging Mass Cytometry (IMC) UMAPs displaying subset pancreas cells from all donors, highlighting the islet-endocrine cells (top) and immune cells (bottom). **(B)** Representative IMC images and spatial reconstructions of islet-endocrine (top) and islet-immune (bottom) fractions. Scale bar = 100μm. **(C)** UMAPs of pancreas cell subpopulations, highlighting endocrine cells (top) and immune subsets (bottom). **(D)** Representative CODEX images and spatial reconstructions for islet immune subpopulations. Scale bar = 2000μm. **(E)** Representative islet for an aged donor, displaying C-peptide (yellow), glucagon (green), CD45 (white) and CD8 (magenta), with inset illustrating alpha-CD8 T cell interaction. Scale bar; image = 100μm, inset = 10μm. **(F-G)** Immune cell proportions within the total pancreatic immune pool, collected by both IMC (F) and CODEX (H) imaging platforms. **(H)** Density plots illustrating networking distance between CD8^+^ T cells distance and alpha cells, median CD8^+^-alpha cell distance, and the percentage of alpha cells localized within ≤50μm of CD8^+^ T cells. **(I)** Density plots showing network distance between CD8⁺ T cells and beta cells, median CD8⁺-beta cell distance, and the percentage of beta cells localized within ≤50μm of CD8⁺ T cells. **(J)** Representative IMC images illustration alpha cells (GCG, red) beta cells (C-peptide, white), and scaled HLA-ABC expression. Scale bar = 100μm. **(K-L).** Quantification of HLA-ABC (K) and HLA-DR (L) intensity from segmented alpha cells and beta cells. IMC, imaging mass cytometry; CyTOF, Cytometry by time-of-flight; ROI, region of interest. Welch’s two-sample *t-test* was used to compare young and old donors in panels F-G, H-I, and K-L. Data are mean ± SEM (panels F-I), median ± SEM (K-L).

This framework enabled spatial mapping of all major pancreatic and immune cell types in both IMC (**Figure 4A-B**) and CODEX (**Figure 4C-D, Supplementary Figure S7A**) platforms to quantify immune and endocrine cell composition and their molecular phenotypes. Using this approach, we found no differences in endocrine islet cell composition (**Supplementary Figure S7B**) and expected age-associated decreases in beta cell identity markers PDX-1, NKX6-1, and PAX6 (**Supplementary Figure S7C**) (*44*). This data confirms loss of beta cell identity during aging without the activation of cell death (*39, 44*). Importantly, we found that ND aging pancreases have significant expansion of CD45^+^CD3^+^ T cells predominantly driven by accumulation of CD8^+^ T-cells that can be found inside the islet (**Figure 4B**, **4D-E, and Supplementary Figure S7D and S8A**). No age-associated differences were detected for CD4^+^ T cells or in total macrophage density (**Figure 4F-G**).

To further investigate the organization of endocrine-immune interactions underlying CD8^+^ T-cell expansion, we utilized spatial coordinates to identify nearest-neighbor cell endocrine beta and alpha cells within a 50μm radius of CD8 T cells, as established previously by other groups (*62–64*). This revealed that older pancreases have a higher fraction of CD8^+^ T cells in the vicinity or in direct contact with alpha cells (**Figure 4E** and **4H, Supplementary Figure S8A**), which was not observed for beta cells (**Figure 4I**).

During an inflammatory event, CD8^+^ T cells engage neighboring cells by interaction between the T cell receptor (TCR) and major histocompatibility complex class 1 (MHC-1) molecules on the interacting cell. Accordingly, the increased spatial proximity between alpha cells and CD8^+^ T cells correlated with a significant increase in the expression of MHC-I complex HLAABC molecules in alpha cells (**Figure 4J**). In fact, older alpha cells express ∼5x more HLA-ABC than neighboring beta cells, and both cell types express near background levels of MHC-II HLADR molecules (**Figure 4L**). Because alpha cell mass was unchanged with aging, this suggests that despite significant interactions between alpha cells and CD8^+^ T cells, aging is not associated with alpha cell loss (as indicated by the single-cell transcriptomics data (**Supplementary Figure S1H**).

Together, these findings demonstrate that human islet and pancreatic aging landscapes are defined by increased pro-inflammatory signaling from alpha cells that correlates with elevated CD8^+^ T-cell density *in situ*. Given that MHC molecules present antigenic peptides to T cells via TCR engagement, these results suggest that alpha cell-intrinsic inflammatory pathways may orchestrate immune activation and islet inflammation during aging.

### T2D expands pancreatic CD8^+^ T cells that shift towards terminal effector-memory phenotypes

To determine how these pancreas and immune aging signatures overlap with T2D pathogenesis, we analyzed CODEX data from T2D donors profiled through HPAP (**Supplementary Table S12**). Cohorts were matched for age and BMI; however, T2D donors displayed expected impairments in glycemia, evidenced by higher HbA1c and reduced C-peptide (**Figure 5A**). Integration of 942,724 single-cell profiles identified and characterized endocrine (99,744 cells) and immune (195,140 cells) compartments (**Figure 5B**). Our endocrine cell clustering approach identified no changes to major islet cell populations and highlighted two distinct beta cell populations: a CPEP^+^CHGA^+^ population and a CPEP^+^CHGA^+^COL4A1^+^COL6^+^ population that was enriched in T2D (**Figure 5C**). These cells exhibited higher interaction with fibrotic extra-cellular matrix (ECM) and thus were detected as being “fibrotic” by our expansion-based cell segmentation algorithms that captured ECM signals from neighboring regions (**Supplementary Figure S8A**). Importantly, T2D beta cells were marked by lower CPEP, CHGA, NKX6-1 expression (**Figure 5D**), while alpha cells had attenuated SOX9 (**Figure 5E**), consistent with compromised endocrine cell identity during T2D (*65*). Within immune compartments, we observed no changes in macrophages densities (**Figure 5F**), and a significant ∼2-fold increase in pancreas CD8^+^ T-cell density (**Figure 5G**). Together, these findings demonstrate that T2D amplifies CD8⁺ T-cell density to mirror patterns observed during aging in ND pancreases.

**Figure 5.**
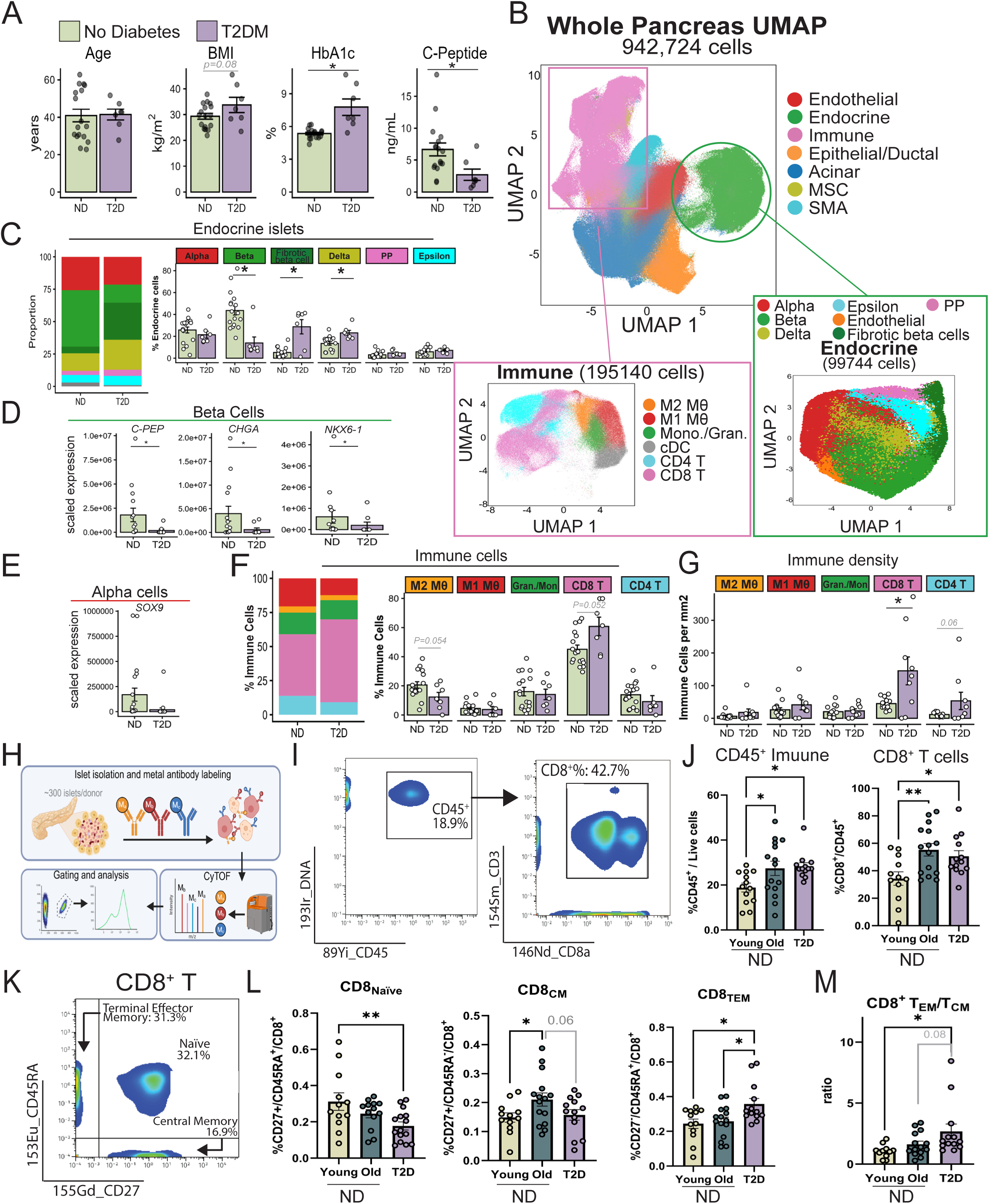
T2D expands pancreatic CD8^+^ T-cell populations and shifts islet CD8^+^ T-cell states from towards terminal effector-memory phenotypes. **(A)** Clinical characteristics ND and T2D adults included for HPAP CODEX imaging. **(B)** UMAP of integrated pancreas samples (942,724 cells) were segmented into immune cells (195,140 cells) and islet endocrine (99,744 cells) populations. **(C)** Identified islet endocrine cell populations, including alpha (GCG^+^), beta (C-PEP^+^), “fibrotic” beta cells (C-PEP^+^, COL4A1^+^, COL6^+^), delta (SST^+^), pancreatic polypeptide (PP^+^), and epsilon (ghrelin^+^) cells. **(D)** Scaled beta cell expression of C-PEP, CHGA, and NKX6-**(E)** Scaled alpha cell expression of SOX9. **(F)** Proportions of major pancreatic immune subsets, including M2-like macrophages (CD163^+^CD68^+^), M1-like macrophages (CD68^+^CD163^-^), granulocytes/monocytes (CD66b^+^), CD8^+^ T cells (CD3^+^CD8^+^), and CD4^+^ T cells (CD3^+^CD4^+^). **(G)** Immune cell density expressed as cells per mm^2^. **(H)** CyTOF analysis of islets isolated from young ND (n=12), old ND(n=15), and T2D (n=13) donors. **(I)** Representative gating of viable CD45^+^ immune cells (left) and CD3^+^CD8^+^ T cells (right), sourced from HPAP-164. **(J)** Proportion of CD45^+^ immune cells among viable islet cells (left) and CD3^+^CD8^+^ within the CD45^+^ compartment (right). **(K)** Representative two-parameter (CD27 x CD45RA) plot illustrating CD8^+^ T-cell differentiation states; (1) naïve CD27^+^CD45RA^+^, (2) central memory (CD27^+^CD45RA^-^), 3) and terminal effector memory (CD27^-^CD45RA^+^). **(L)** Distribution of CD8^+^ T cells across naïve CD27^+^CD45RA^+^; CD8_naïve_), central memory (CD27^+^CD45RA^-^; CD8_CM_), and terminal effector memory (CD27^-^CD45RA^+^; CD8_TEM_) phenotypes. **(M)** CD8^+^ T_EM_/T_CM_ ratio, representing the relative expansion of terminal effector-memory CD8^+^ T cells compared with the central memory pool. Welch’s two sample *t-test* was used for comparisons between pool ND and T2D donors in panels A and C-G. One-way *ANOVA* compared young, old and T2D donors in panels J and L-M. DC, dendritic cell; cDC1, conventional cell; PP, pancreatic polypeptide; NK, natural killer. *p< 0.05, **p<0.01 for age (old vs. young) and ND vs. T2D groups. Data are mean ± SEM.

To resolve whether the increased adaptive immune phenotype observed in both aging and T2D pancreases was localized within islets, we analyzed HPAP cytometry by time-of-flight (CyTOF) data from isolated islets of young (26±8 years, n=13), old (54±5 years, n=14), and T2D (50±8 years, n=13) human donors (**Figure 5H, Supplementary Table S13-14**). Both aging and T2D presented greater CD45^+^ live immune cell abundance compared with young adults, corresponding with higher frequencies of islet CD8^+^ T cells within the CD45^+^ immune compartment (**Figure 5I-J, Supplementary Figure S9**). CD8^+^ T cells differentiate along a spectrum from naïve (CD8_naïve_) to CD8_CM_ and CD8_TEM_ states in response to TCR activation by antigen and cytokine stimulation. In T2D islets, we observed a reduction in CD45RA^+^CD27^+^ CD8_naïve_ T cells compared to young adults, whereas aging increased CD45RA^-^CD27^+^ CD8_CM_ T cells (**Figure 5K-L**). T2D islets further shifted CD8^+^ T-cell populations towards a higher CD45RA^+^CD27^-^ CD8_TEM_ phenotype, resulting in elevated effector-to-central memory ratio that was not observed in aging ND samples (**Figure 5L-M**).

Together, these data show that T2D intensifies pancreas and islet CD8⁺ T-cell infiltration patterns found during aging to drive their progression towards terminal effector-memory states.

### Calorie restriction improves glucose tolerance and dampens alpha cell inflammation in mice

CR is a dietary intervention approach that can normalize glycemia during prediabetes and T2D, likely by delaying the onset of workload stress and aging hallmarks in beta cells (*32, 66–69*). We therefore tested whether CR could reverse aging signatures in alpha and beta cells *in vivo*. To achieve this, 72-week-old C57BL/6J male mice were subjected to a 2-month, 20% CR feeding regimen, while age-matched controls remained on ad-libitum (AL) diet (**Figure 6A**). As expected, CR mice consumed 80% of total kcal intake compared to AL-fed controls (**Supplementary Figure S10A-B**). This intervention induced a steady decline in body weight over 2 months, resulting in ∼12% weight loss (**Figure 6B, Supplementary Figure S10C-D**) predominantly due to a loss in fat mass (**Supplementary Figure S10E-F**). Of note, CR mice also experienced reductions in liver and epididymal white adipose tissue (eWAT) mass due to reduced lipid droplet size (**Supplementary Figure S10G-J**).

**Figure 6.**
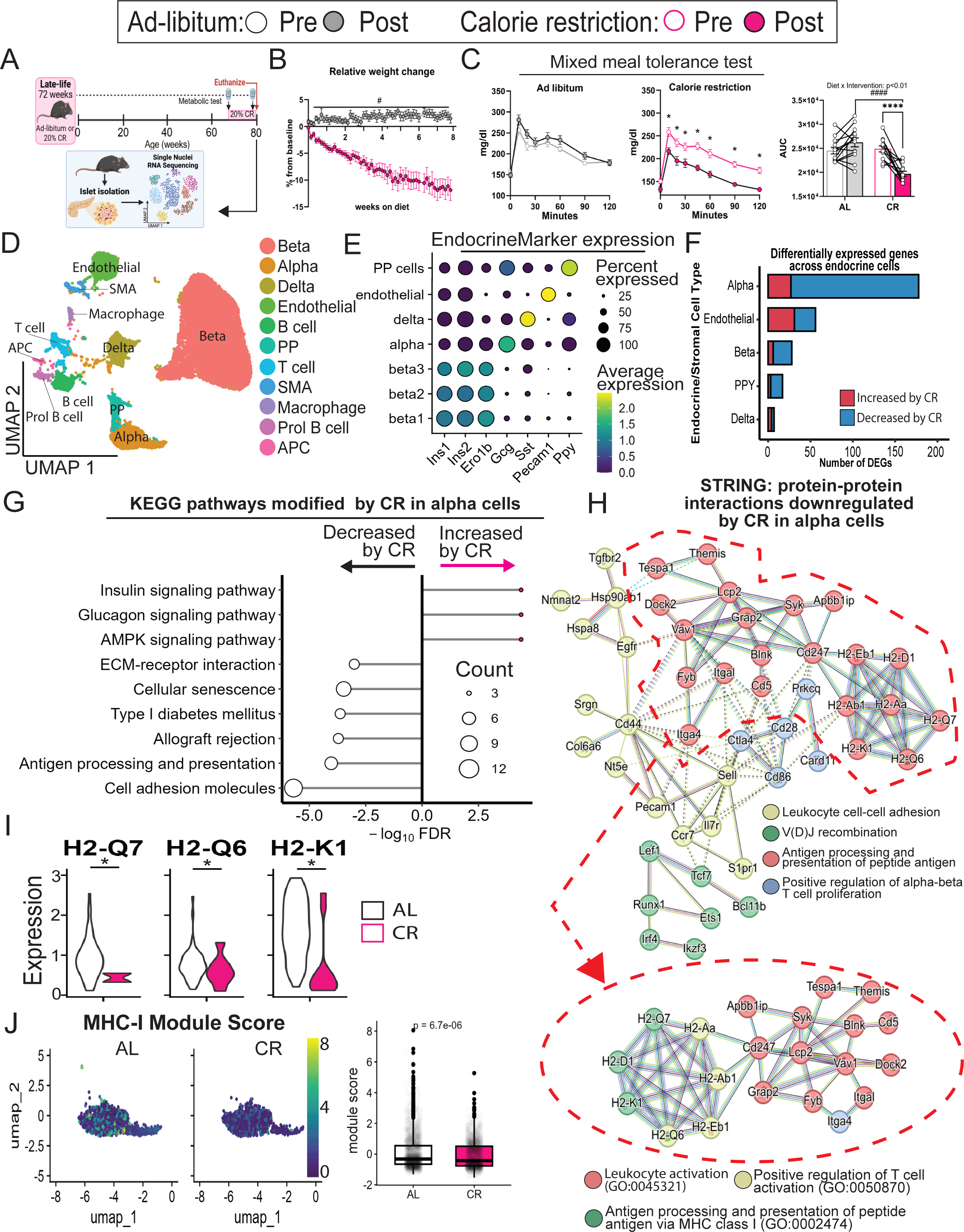
Late-life CR improves glucose tolerance while downregulating alpha cell TCR activation and antigen presentation and processing. **(A)** 72-week-old C57Bl/6J mice were fed ad-libitum (AL, control) or subjected to 20% calorie restriction (CR) for 2 months. Islets were collected at 80 weeks and processed for IHC and snRNA-seq. **(B)** Percent change in body weight relative to baseline. **(C)** MTTs were performed at baseline (72 weeks) and after the intervention (80 weeks). AUC is presented for each test before and after dietary intervention. **(D)** UMAP of 32,380 islet cells. **(E)** Selected marker genes for endocrine islet subpopulations by scaled gene expression and percent cell expression. **(F)** Number of DEGs affected by CR in endocrine/endothelial cell populations. **(G)** KEGG pathways significantly modified by CR in alpha cells. **(H)** Protein-protein interaction networks predicted from genes downregulated by CR in alpha cells, and subset by ‘leukocyte activation’ subcluster in red. **(I)** Scaled single-cell expression of MHC class I in alpha cells. **(J)** Scaled expression UMAP and box plot showing relative MHC-I module scores, derived from *H2-K1, H2-D1, H2-Q2, H2-Q5, H2-Q6, H2-Q7, and H2-Q12* genes, in AL and CR alpha cells. AUC, area under curve; IHC, immunohistochemistry; MTT, mixed meal tolerance test; snRNA-seq, single nucleus RNA sequencing. #p< 0.05 for dietary groups (AL vs. CR); ****p<0.0001 for intervention (Pre- vs. Post-CR).

Metabolic phenotyping and behavioral analysis revealed comparable overall energy expenditure and oxygen consumption between AL and CR mice (**Supplementary Figure S10KN**). However, carbon dioxide production and carbohydrate oxidation were higher in CR mice during the dark period when food was available (**Supplementary Figure S10O-R**), indicating a higher preference for carbohydrate metabolism in CR mice during active feeding periods and a rapid shift towards fatty acid metabolism during inactivity and food withdrawal. To assess how CR influences glucose homeostasis in aged mice, we performed mixed meal tolerance tests (MTTs; 2 g glucose/kg body weight). Baseline MTTs were conducted at 72 weeks, prior to CR. After 2 months of CR, old mice exhibited significantly improved glucose tolerance at 80 weeks, attributed to enhanced insulin sensitivity (**Figure 6C**). This improvement was accompanied by ∼50% reductions in fasting and glucose-stimulated beta cell insulin secretion (**Supplementary Figure S10S**). This indicates that CR preserves aging beta cell function by reducing the relative insulin secretory demand required to maintain normoglycemia, demonstrating that known CR-mediated functional beta cell adaptation (*33*) remains intact in aging (**Supplementary Figure S10T**).

We previously showed that CR modulates beta cell heterogeneity by activating pro-longevity and pro-homeostasis transcriptional mechanisms that enhance beta cell protein and organelle quality-control mechanisms (*33*). This transcriptional response is driven by the activation of TF-directed GRNs that increase the expression of beta cell identity and longevity genes (*33*). Therefore, to determine whether CR elicits similar adaptative programs to old beta cells during CR, we performed single nucleus RNA sequencing (snRNA-seq, 10X genomics) on islet nuclei isolated from AL and CR-fed mice (**Figure 6A**). A total of 32,380 cells were sequenced (AL=17,344, CR=15,306) to identify all major islet cell types, including beta cells (*Ins1*, *Ins2*, and *Ero1b*), alpha cells (*Gcg*), delta cells (*Sst*), endothelial cells (*Pecam1*), and pancreatic polypeptide cells (*Ppy*) (**Figure 6D-E, Supplementary Figure S11A**). Approximately 65% of the sequenced cells were beta cells, and CR did not impact the relative composition of the heterogeneous beta cell population or islet cytoarchitecture composition (**Supplementary Figure S11B-D**). Using pseudobulk analysis of beta cells followed by KEGG and STRING databases, we observed a narrow range of CR-mediated effects on beta cell gene expression, limited to downregulation of few ER protein processing and folding (*Hspa1a/b/HSP70, Hspa8, Hsp90ab1, Dnaja1/HSP40, and Calr)*, cell stress (*Ypel2*, *Txnip*), and insulin synthesis (*Ins1*) genes (**Figure 6F and Supplementary Figure S11E-J**). These data indicate that CR has a very limited impact on beta cell transcriptional programs in aged mice despite a significant downregulation of insulin release *in vivo*.

Remarkably, CR exerted a stronger transcriptional effect on alpha cells, modulating nearly five times more genes than beta cells (184 vs. 36 differentially expressed genes; **Figure 6F, Supplementary Table S15**). Differential gene expression and pathway enrichment analyses of old CR alpha cell gene signatures revealed upregulation of pathways associated with glucagon, glucose and fatty acid metabolism (*Gcg, Gys2, Acacb*, and *Pgc1a*/*Ppargc1a)* (**Figure 6G**). In contrast, CR suppressed 159 genes involved in TCR signaling, antigen presentation, and immune cell recruitment pathways (**Figure 6H, Supplement Figure S12**). This included genes involved in cell-cell signaling via MHC class I (H2-D1, *H2-Q7*, *H2-Q6*, *H2-K1*) and some MHC class II (*H2-Aa*, *H2-Ab1*) antigens **(Figure 6H-I)**. Because MHC-I antigens is more prominent in islet endocrine cells than MHC-II, we constructed an MHC-I module score based on heavy and light chain genes (*H2-K1, H2-D1, H2-Q2, H2-Q5,H2-Q6,H2-Q7,H2-Q12*) detected in our scRNA-seq data, and found the module score was reduced by CR (**Figure 6J**).

Taken together, these findings demonstrate that 2 months of CR reduces pro-inflammatory signaling in aging alpha cells in mice, and it induces a state of beta cell rest that does not involve significant changes to beta cell transcriptional mechanisms.

### CR suppresses alpha cell immune signaling to reduce islet inflammation

To further investigate how CR-mediated modulation of aging alpha cell inflammatory gene expression influences the cellular and molecular profile of islet immune cells, we quantified the relative composition and transcriptional signatures immune cell types identified by snRNA-seq (**Figure 7A**).

**Figure 7.**
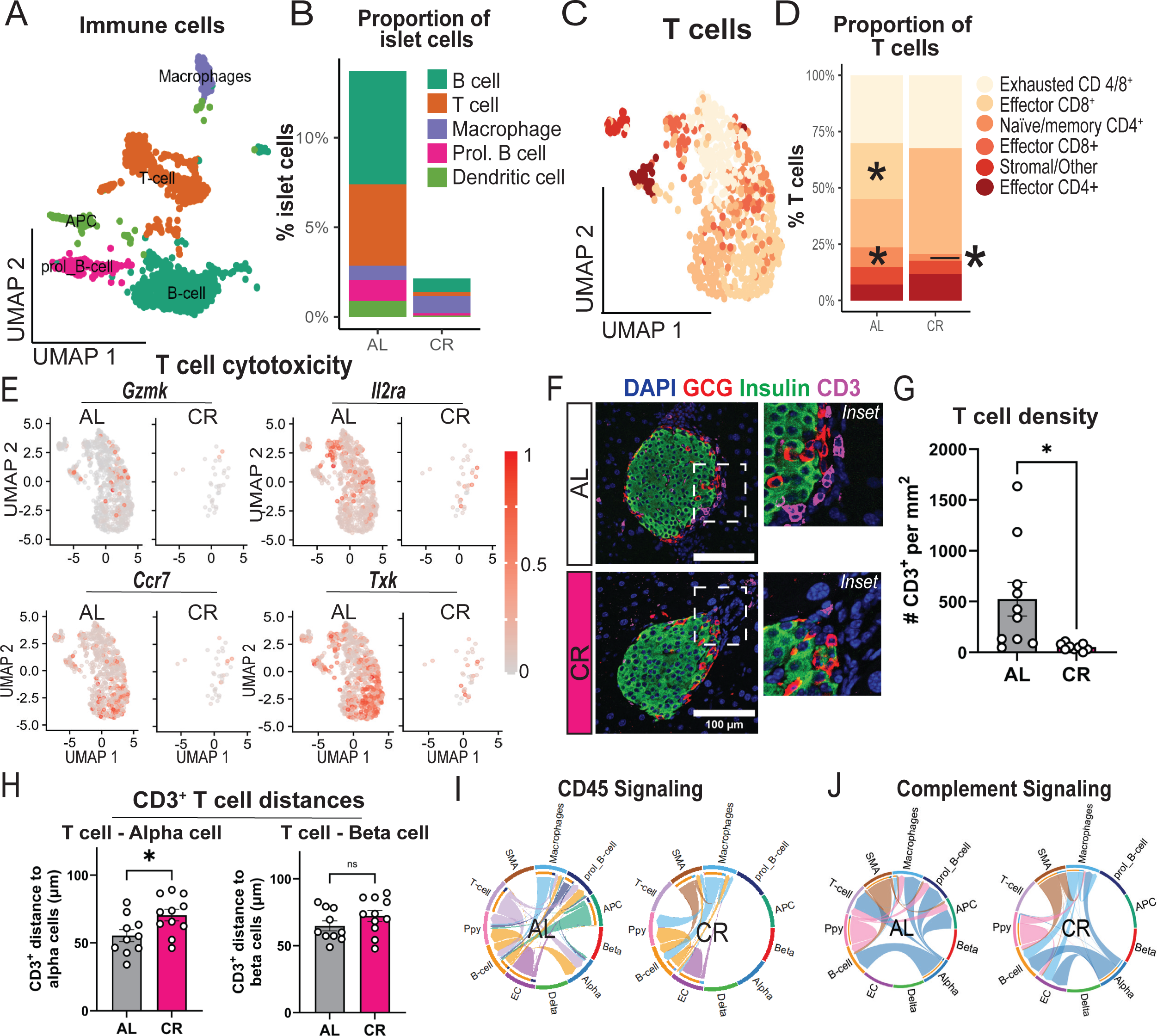
Late-life CR attenuates islet adaptive CD8^+^ effector phenotypes and reduces alpha cell-mediated T-cell recruitment. **(A)** UMAP of subset immune cell populations (B cells, T cells, macrophages, proliferating B cells, and APCs [dendritic cells] from AL and CR groups. **(B)** Proportion of immune cell populations relative to total islet cells. **(C)** UMAP of T-cell populations from AL and CR groups. **(D)** Proportion of total T cells; * denotes effector CD8^+^ T cells. **(E)** Expression of Th1-associated genes: *Gzmk*, *Cd25/Il2ra*, *CCr7*, and *Txk*. **(F)** Representative IHC staining for alpha cells (glucagon, red), beta cells (insulin, green) and T cells (CD3, purple). Scale bar = 100μm. **(G)** Pancreatic T-cell density (CD3^+^ T-cell counts per mm^2^). **(H)** T-cell-endocrine cell networks, quantifying the average distance of CD3^+^ T cells to alpha cells (left), and beta cells (right). **(I-J)** Intercellular communication networks showing inferred CD45 signaling and complement signaling interactions, respectively.

This revealed a significant decrease in B cells, T cells, proliferating B cells, and antigen presenting cells (APCs) in CR mouse islets, whereas the relative density and transcriptional profile of islet-resident macrophages were unchanged (**Figure 7B**). Old CR mouse islets had a marked reduction in islet T cells, including a near total ablation of cytotoxic CD8^+^ effector T cells (**Figure 7C-D**). In addition, the remaining CD8^+^ T cells in CR islets had reduced expression of effector markers granzyme K (*Gzmk*), IL-2 receptor (Il2ra/*Cd25*), Ccr7, and *Txk*; all markers of activated CD8^+^ T cells (**Figure 7E**). We validated these changes *in situ* using immunohistochemistry and confocal microscopy of AL and CR old pancreases, followed by spatial mapping of alpha cells, beta cells, and CD3^+^ T cells (**Figure 7F)**. This approach revealed a significant reduction in T-cell density (**Figure 7G**), and increased distance between T cells and alpha cells (**Figure 7H**). The proportion of T cells in proximity (within 6μm) to alpha cells trended higher (p=0.07), whereas this enrichment was not observed for beta cells (**Supplementary Figure S13A-B**). Total macrophage content in proximity to islets was not altered by CR (**Supplementary Figure 13C-E**), and the relative proportions of pro-inflammatory (M1-like) and anti-inflammatory (M2-like) macrophages were likewise unchanged (**Supplementary Figure S13F-K**). These data support recent findings of immune cell infiltration in the aging mouse islet (*70*) and demonstrate that CR modulates the recruitment of CD8^+^ T cells to the islet by reducing their preferential proximity to old alpha cells.

To identify communication pathways linking aged alpha cells with CD8^+^ T cells and their modulation by CR, we performed receptor-ligand inference analysis using CellChat (*71, 72*) to reconstruct cell-cell communication networks in islets from AL and CR mice (*73*). This analysis revealed a downregulation of intercellular signaling between islet immune cells and endocrine cell types, including alpha, beta, and delta cells (**Supplementary Figure S14A**). This was underscored by a significant decrease in CD45-based interactions between old alpha cells and T- and B-cells in CR mice (**Figure 7I**), accompanied by reduced complement signaling, particularly toward T cells (**Figure 7J**). Moreover, CR attenuated T cell co-stimulation, as evidenced by reduced CD80 signaling from APCs and reduced CD86 signaling from B cells to T cells (**Supplementary Figure S14B–C**). Chemokine signaling via CCL and CXCL pathways towards T cells was similarly reduced under CR (**Supplementary Figure S14D-E**).

Overall, these results indicate that CR profoundly remodels the aging islet immune environment by suppressing alpha cell inflammatory and MHC-I signaling, thereby reducing CD8⁺ T cell recruitment and cytotoxicity, and diminishing intercellular communication between alpha cells and adaptive immune populations.

## Discussion

Aging is associated with an active and low-grade inflammatory process in different tissues, where inflammation is a key signature of several age-associated degenerative and metabolic diseases, including Alzheimer’s (*74*), and obesity and diabetes (*75*). In the pancreas, aging is associated with increased islet macrophage infiltration and vascular fibrosis (*38*), and share other key hallmarks of T1D and T2D islet pathophysiology such as beta cell ER stress, compromised beta cell identity and function, and beta cell senescence (*15, 38, 39, 43–45, 56, 57, 76–82*). Importantly, recent studies have implicated the activation of inflammatory responses in human alpha cells during T1D and T2D (*83, 84*).

Here, we identified the activation of immune signaling and inflammation as a key signature of aging human islets from non-diabetic individuals. This phenotype is defined by active IFNγ signaling and MHC-I expression in old alpha cells due to activation of IFNγ-responsive TFs IRF1, NFKB1, and STAT1. Activation of alpha cell immune signaling during aging translates into increased physical interactions between old alpha cells with pancreas CD8^+^ T cells, which accumulate in the human islet microenvironment during aging. Strikingly, most old islet CD8^+^ cells have a “memory-like” phenotype (CD27^+^), thus indicating that these cells have been exposed to T cell antigens. While our study does not identify which antigen is presented by alpha cell MHC-Is, a recent study where human islets were exposed to cytokine stress (e.g., IFNγ) identified several new MHC-I antigens, many of which are enriched in aging alpha cells (i.e., NFKB1, IRF9, TAP1, IFIT3, PSMB8, and STAT1)(*76*).

This aging alpha cell inflammatory phenotype correlates with an increase in the expression of several AA catabolism enzymes. This could explain the observed increase in amino-acid-dependent glucagon release from isolated human islets and the elevated levels of glucagon in old ND humans after a meal (*2*). Notably, elevated glucagon secretion after a mixed meal and AA infusions is also observed in adults with T2D (*19–21*), thus suggesting that alpha cell inflammation could also play a role during T2D pathogenesis. In fact, while we did not observe changes in the density of islet CD8+ cells between ND and T2D islets, most T2D islet CD8^+^ cells have an effector-like signature (CD45RA^+^CD8^+^). This suggests that infiltration of CD8^+^ cells in aging human islets occurs independently of T2D and further suggests that islet inflammation in individuals with T2D onset during old age involves activation of alpha cell-interacting and “resident” CD8^+^ cells and broad islet inflammation signatures (*38, 80*).

In species spanning from yeast to humans, CR has been shown to extend lifespan and health span (*32, 68, 69, 85–87*). The benefits of CR depend on improvements in glucose homeostasis, enhanced insulin sensitivity, and reduced metabolic demand for insulin release from beta cells leading to beta cell rest (*33, 69, 88, 89*). In addition, CR can reduce overall inflammation signatures in different models, including humans (*32*). Our experiments with 80-week-old mice on CR for 2 months demonstrate that the insulin-sensitizing and beta cell functional adaptation pathways promoted by CR are not impaired during aging and that these adaptations do not require significant transcriptional reorganization. This is likely explained by the limited plasticity of aging beta cells to respond to metabolic challenges (*90*), however the mechanisms underlying this limited plasticity remain unknown. Notably, a recent study has shown that aging of mouse endocrine islet cells is linked to immune activation and MHC-I gene expression (*82*).

While we did not observe this signature in aging beta cells and delta cells, CR acted on old alpha cells to significantly suppress the expression of pro-inflammatory genes, including MHC-I signaling pathways. This CR-dependent modulation dampened CD45-mediated communication between immune cells and alpha cells, which was sufficient to reduce the density of T cells in aging islet periphery.

In conclusion, our findings identify the relationship between alpha cells and immune cells within the islet of Langerhans during aging that may underlie pro-inflammatory injury and promote T2D development during old age.

## Data availability

All source data and metadata, including IMC, CODEX, snRNA-seq, and source code data are publicly available and can be downloaded from Zenodo (10.5281/zenodo.17874622). Human islet single cell RNAseq uniformly reprocessed is available via PanKbase Zenodo (https://zenodo.org/records/15596314). Raw FASTQ data from mouse islet sequencing data are deposited at Gene Expression Omnibus and processed data is available on Github and Zenodo.

## Acknowledgements

We acknowledge the following Vanderbilt University (VU) and Vanderbilt University Medical Center (VUMC) core facilities: VUMC Hormone Assay & Analytical Services Core (NIH DK020593), VU Metabolic Mouse Phenotyping Center [VMMPC (NIH DK059637; www.vmmpc.org)], Translational Pathology Shared Resource (NCI/NIH Cancer Center Support Grant 5P30 CA68485-19), and Vanderbilt Technologies for Advanced Genomics (VANTAGE) core laboratory. VANTAGE is supported in part by Clinical and Translational Science Award Grant 5UL1 RR024975-03, Vanderbilt Ingram Cancer Center Grant P30 CA68485, Vanderbilt Vision Center Grant P30 EY08126, and National Institutes of Health/National Center for Research Resources Grant G20 RR030956. This manuscript used data acquired from the database (https://hpap.pmacs.upenn.edu/) of the Human Pancreas Analysis Program (HPAP; RRID:SCR_016202)(*60, 61*). HPAP is part of a Human Islet Research Network (RRID:SCR_014393) consortium (UC4-DK112217, U01-DK123594, UC4-DK112232, and U01-DK123716). This work includes data and/or analyses from HumanIslets.com funded by the Canadian Institutes of Health Research, JDRF Canada, and Diabetes Canada (5-SRA-2021-1149-S-B/TG 179092) with data from islets isolated by the Alberta Diabetes Institute IsletCore with the support of the Human Organ Procurement and Exchange program, Trillium Gift of Life Network, BC Transplant, Quebec Transplant, and other Canadian organ procurement organizations with written informed donor consent as approved by the Human Research Ethics Board at the University of Alberta (Pro00013094). we would like to acknowledge Dr Patrick E MacDonald from ADI – Edmonton (Canada) for critical comments and inputs during data collection and manuscript writing stages of this project. We would also like to acknowledge NIH grant R01AG040178.

## Author Contributions

M.S., S.S., JP.C., and R.A.D designed experiments. M.S., A.C., M.C., G.F., S.S., JP.C., A.H., C.E., A.P., and R.A.D. collected data and assisted with data analysis and project management. M.S., S.S., JP.C., and R.A.D. completed data analysis. M.S. and R.A.D. interpreted data and wrote the manuscript.

## REFERENCES

1. National Diabetes Statistics Report. Centers for Disease Control and Prevention, (2024).

2. R. Basu et al., Mechanisms of the age-associated deterioration in glucose tolerance: contribution of alterations in insulin secretion, action, and clearance. Diabetes 52, 1738–1748 (2003).

3. K. F. Petersen et al., Mitochondrial dysfunction in the elderly: possible role in insulin resistance. Science 300, 1140–1142 (2003).

4. C. C. Cowie et al., Prevalence of diabetes and impaired fasting glucose in adults in the U.S. population: National Health And Nutrition Examination Survey 1999-2002. Diabetes Care 29, 1263–1268 (2006).

5. A. Alpert et al., A clinically meaningful metric of immune age derived from high-dimensional longitudinal monitoring. Nat Med 25, 487–495 (2019).

6. C. Sportès et al., Administration of rhIL-7 in humans increases in vivo TCR repertoire diversity by preferential expansion of naive T cell subsets. J Exp Med 205, 1701–1714 (2008).

7. M. L. Pekalski et al., Postthymic expansion in human CD4 naive T cells defined by expression of functional high-affinity IL-2 receptors. J Immunol 190, 2554–2566 (2013).

8. X. Sun et al., Longitudinal analysis reveals age-related changes in the T cell receptor repertoire of human T cell subsets. J Clin Invest 132, (2022).

9. O. Cabrera et al., The unique cytoarchitecture of human pancreatic islets has implications for islet cell function. Proceedings of the National Academy of Sciences 103, 2334–2339 (2006).

10. M. Cnop et al., The long lifespan and low turnover of human islet beta cells estimated by mathematical modelling of lipofuscin accumulation. Diabetologia 53, 321–330 (2010).

11. R. A. E. Drigo et al., Age Mosaicism across Multiple Scales in Adult Tissues. Cell Metabolism 30, 343-+ (2019).

12. S. Perl et al., Significant human beta-cell turnover is limited to the first three decades of life as determined by in vivo thymidine analog incorporation and radiocarbon dating. J Clin Endocrinol Metab 95, E234–239 (2010).

13. C. López-Otín, M. A. Blasco, L. Partridge, M. Serrano, G. Kroemer, Hallmarks of aging: An expanding universe. Cell 186, 243–278 (2023).

14. S. Shrestha et al., Aging compromises human islet beta cell function and identity by decreasing transcription factor activity and inducing ER stress. Science Advances 8, eabo3932 (2022).

15. I. Sandovici et al., Ageing is associated with molecular signatures of inflammation and type 2 diabetes in rat pancreatic islets. Diabetologia 59, 502–511 (2016).

16. L. Li et al., Defects in β-cell Ca2+ dynamics in age-induced diabetes. Diabetes 63, 4100–4114 (2014).

17. G. M. Reaven, Y. D. Chen, A. Golay, A. L. Swislocki, J. B. Jaspan, Documentation of hyperglucagonemia throughout the day in nonobese and obese patients with noninsulin-dependent diabetes mellitus. J Clin Endocrinol Metab 64, 106–110 (1987).

18. K. Færch et al., Insulin Resistance Is Accompanied by Increased Fasting Glucagon and Delayed Glucagon Suppression in Individuals With Normal and Impaired Glucose Regulation. Diabetes 65, 3473–3481 (2016).

19. R. H. Unger, E. Aguilar-Parada, W. A. Müller, A. M. Eisentraut, Studies of pancreatic alpha cell function in normal and diabetic subjects. J Clin Invest 49, 837–848 (1970).

20. D. D’Alessio, The role of dysregulated glucagon secretion in type 2 diabetes. Diabetes Obes Metab 13 Suppl 1, 126–132 (2011).

21. P. Raskin, I. Aydin, T. Yamamoto, R. H. Unger, Abnormal alpha cell function in human diabetes: the response to oral protein. Am J Med 64, 988–997 (1978).

22. P. Raskin, I. Aydin, R. H. Unger, Effect of insulin on the exaggerated glucagon response to arginine stimulation in diabetes mellitus. Diabetes 25, 227–229 (1976).

23. L. Fontana, L. Partridge, V. D. Longo, Extending healthy life span--from yeast to humans. Science 328, 321–326 (2010).

24. J. A. Mattison et al., Caloric restriction improves health and survival of rhesus monkeys. Nat Commun 8, 14063 (2017).

25. C. L. Green, D. W. Lamming, L. Fontana, Molecular mechanisms of dietary restriction promoting health and longevity. Nat Rev Mol Cell Biol 23, 56–73 (2022).

26. R. M. Anderson, D. G. Le Couteur, R. de Cabo, Caloric Restriction Research: New Perspectives on the Biology of Aging. J Gerontol A Biol Sci Med Sci 73, 1–3 (2017).

27. K. Mao et al., Late-life targeting of the IGF-1 receptor improves healthspan and lifespan in female mice. Nat Commun 9, 2394 (2018).

28. G. Y. Liu, D. M. Sabatini, mTOR at the nexus of nutrition, growth, ageing and disease. Nat Rev Mol Cell Biol 21, 183–203 (2020).

29. D. Il’yasova et al., Effects of 2 years of caloric restriction on oxidative status assessed by urinary F2-isoprostanes: The CALERIE 2 randomized clinical trial. Aging Cell 17, (2018).

30. L. Yang, P. Li, S. Fu, E. S. Calay, G. S. Hotamisligil, Defective hepatic autophagy in obesity promotes ER stress and causes insulin resistance. Cell Metab 11, 467–478 (2010).

31. J. O. Pyo et al., Overexpression of Atg5 in mice activates autophagy and extends lifespan. Nat Commun 4, 2300 (2013).

32. O. Spadaro et al., Caloric restriction in humans reveals immunometabolic regulators of health span. Science 375, 671-+ (2022).

33. C. dos Santos et al., Calorie restriction increases insulin sensitivity to promote beta cell homeostasis and longevity in mice. Nature Communications 15, 9063 (2024).

34. L. Matta et al., Chronic intermittent fasting impairs β cell maturation and function in adolescent mice. Cell Rep 44, 115225 (2025).

35. S. Patel, Z. Yan, M. S. Remedi, Intermittent fasting protects β-cell identity and function in a type-2 diabetes model. Metabolism 153, 155813 (2024).

36. M. R. Brown et al., Time-restricted feeding prevents deleterious metabolic effects of circadian disruption through epigenetic control of β cell function. Sci Adv 7, eabg6856 (2021).

37. L. Mateus Gonçalves, E. Pereira, J. P. Werneck de Castro, E. Bernal-Mizrachi, J. Almaça, Islet pericytes convert into profibrotic myofibroblasts in a mouse model of islet vascular fibrosis. Diabetologia 63, 1564–1575 (2020).

38. J. Almaca et al., Young capillary vessels rejuvenate aged pancreatic islets. Proceedings of the National Academy of Sciences of the United States of America 111, 17612–17617 (2014).

39. J. J. Wright et al., Exocrine Pancreas in Type 1 and Type 2 Diabetes: Different Patterns of Fibrosis, Metaplasia, Angiopathy, and Adiposity. Diabetes 73, 1140–1152 (2024).

40. J. D. Ewald et al., HumanIslets.com: Improving accessibility, integration, and usability of human research islet data. Cell Metab 37, 7–11 (2025).

41. J. Kolic et al., Proteomic predictors of individualized nutrient-specific insulin secretion in health and disease. Cell Metab 36, 1619–1633.e1615 (2024).

42. J. Camunas-Soler et al., Patch-Seq Links Single-Cell Transcriptomes to Human Islet Dysfunction in Diabetes. Cell Metabolism 31, 1017-+ (2020).

43. C. Aguayo-Mazzucato et al., Acceleration of beta Cell Aging Determines Diabetes and Senolysis Improves Disease Outcomes. Cell Metabolism 30, 129-+ (2019).

44. S. Shrestha et al., Aging compromises human islet beta cell function and identity by decreasing transcription factor activity and inducing ER stress. Science Advances 8, 17 (2022).

45. M. Enge et al., Single-Cell Analysis of Human Pancreas Reveals Transcriptional Signatures of Aging and Somatic Mutation Patterns. Cell 171, 321–330.e314 (2017).

46. S. Tritschler, F. J. Theis, H. Licked, A. Bottcher, Systematic single-cell analysis provides new insights into heterogeneity and plasticity of the pancreas. Molecular Metabolism 6, 974–990 (2017).

47. P. Augsornworawat, K. G. Maxwell, L. Velazco-Cruz, J. R. Millman, Single-Cell Transcriptome Profiling Reveals β Cell Maturation in Stem Cell-Derived Islets after Transplantation. Cell Rep 32, 108067 (2020).

48. J. Chiou et al., Single-cell chromatin accessibility identifies pancreatic islet cell type- and state-specific regulatory programs of diabetes risk. Nature Genetics 53, 455-+ (2021).

49. N. C. Bramswig et al., Epigenomic plasticity enables human pancreatic α to β cell reprogramming. J Clin Invest 123, 1275–1284 (2013).

50. D. Avrahami et al., Aging-Dependent Demethylation of Regulatory Elements Correlates with Chromatin State and Improved β Cell Function. Cell Metabolism 22, 619–632 (2015).

51. D. Avrahami et al., Single-cell transcriptomics of human islet ontogeny defines the molecular basis of β-cell dedifferentiation in T2D. Mol Metab 42, 101057 (2020).

52. C. Su et al., 3D chromatin maps of the human pancreas reveal lineage-specific regulatory architecture of T2D risk. Cell Metabolism 34, 1394-+ (2022).

53. H. E. Arda et al., Age-Dependent Pancreatic Gene Regulation Reveals Mechanisms Governing Human beta Cell Function. Cell Metabolism 23, 909–920 (2016).

54. M. Fasolino et al., Single-cell multi-omics analysis of human pancreatic islets reveals novel cellular states in type 1 diabetes. Nature Metabolism 4, 284-+ (2022).

55. A. R. Patil et al., Single-cell expression profiling of islets generated by the Human Pancreas Analysis Program. Nat Metab 5, 713–715 (2023).

56. C. Aguayo-Mazzucato et al., beta Cell Aging Markers Have Heterogeneous Distribution and Are Induced by Insulin Resistance. Cell Metabolism 25, 898-+ (2017).

57. P. J. Thompson et al., Targeted Elimination of Senescent Beta Cells Prevents Type 1 Diabetes. Cell Metab 29, 1045–1060.e1010 (2019).

58. B. van de Sande et al., A scalable SCENIC workflow for single-cell gene regulatory network analysis. Nature Protocols 15, 2247–2276 (2020).

59. A. N. Schep, B. Wu, J. D. Buenrostro, W. J. Greenleaf, chromVAR: inferring transcription-factor-associated accessibility from single-cell epigenomic data. Nat Methods 14, 975–978 (2017).

60. K. H. Kaestner, A. C. Powers, A. Naji, M. A. Atkinson, NIH Initiative to Improve Understanding of the Pancreas, Islet, and Autoimmunity in Type 1 Diabetes: The Human Pancreas Analysis Program (HPAP). Diabetes 68, 1394–1402 (2019).

61. S. N. Shapira, A. Naji, M. A. Atkinson, A. C. Powers, K. H. Kaestner, Understanding islet dysfunction in type 2 diabetes through multidimensional pancreatic phenotyping: The Human Pancreas Analysis Program. Cell Metab 34, 1906–1913 (2022).

62. É. Korpos et al., Identification and characterisation of tertiary lymphoid organs in human type 1 diabetes. Diabetologia 64, 1626–1641 (2021).

63. M. H. Wu et al., Single-cell analysis of the human pancreas in type 2 diabetes using multi-spectral imaging mass cytometry. Cell Reports 37, 15 (2021).

64. Y. J. Wang et al., Multiplexed In Situ Imaging Mass Cytometry Analysis of the Human Endocrine Pancreas and Immune System in Type 1 Diabetes. Cell Metab 29, 769–783.e764 (2019).

65. C. Talchai, S. H. Xuan, H. V. Lin, L. Sussel, D. Accili, Pancreatic beta Cell Dedifferentiation as a Mechanism of Diabetic beta Cell Failure. Cell 150, 1223–1234 (2012).

66. C. Dos Santos et al., Caloric restriction promotes beta cell longevity and delays aging and senescence by enhancing cell identity and homeostasis mechanisms. Res Sq, (2023).

67. E. Duregon et al., Prolonged fasting times reap greater geroprotective effects when combined with caloric restriction in adult female mice. Cell Metab 35, 1179–1194.e1175 (2023).

68. S. J. Mitchell et al., Effects of Sex, Strain, and Energy Intake on Hallmarks of Aging in Mice. Cell Metabolism 23, 1093–1112 (2016).

69. H. H. Pak et al., Fasting drives the metabolic, molecular and geroprotective effects of a calorie-restricted diet in mice. Nature Metabolism 3, 1327-+ (2021).

70. S. Jin et al., Inference and analysis of cell-cell communication using CellChat. Nat Commun 12, 1088 (2021).

71. A. Coomans de Brachène et al., Exercise as a non-pharmacological intervention to protect pancreatic beta cells in individuals with type 1 and type 2 diabetes. Diabetologia 66, 450–460 (2023).

72. P. Carapeto et al., Exercise activates AMPK in mouse and human pancreatic islets to decrease senescence. Nature Metabolism, (2024).

73. H. Al Jobori et al., Empagliflozin Treatment Is Associated With Improved β-Cell Function in Type 2 Diabetes Mellitus. J Clin Endocrinol Metab 103, 1402–1407 (2018).

74. C. M. Kellogg et al., Microglial MHC-I induction with aging and Alzheimer’s is conserved in mouse models and humans. Geroscience 45, 3019–3043 (2023).

75. S. SantaCruz-Calvo et al., Adaptive immune cells shape obesity-associated type 2 diabetes mellitus and less prominent comorbidities. Nat Rev Endocrinol 18, 23–42 (2022).

76. P. P. Nanaware et al., The antigen presentation landscape of cytokine-stressed human pancreatic islets. Cell Rep 44, 115927 (2025).

77. L. Marselli et al., beta-Cell inflammation in human type 2 diabetes and the role of autophagy. Diabetes Obesity & Metabolism 15, 130–136 (2013).

78. J. Jelleschitz et al., Insulitis and aging: Immune cell dynamics in Langerhans islets. Redox Biol 82, 103587 (2025).

79. M. De Burghgrave et al., Pancreatic Islet Cells Response to IFNγ Relies on Their Spatial Location within an Islet. Cells 12, (2022).

80. D. Avrahami et al., Single-cell transcriptomics of human islet ontogeny defines the molecular basis of beta-cell dedifferentiation in T2D. Molecular Metabolism 42, 14 (2020).

81. E. Horwitz et al., β-Cell DNA Damage Response Promotes Islet Inflammation in Type 1 Diabetes. Diabetes 67, 2305–2318 (2018).

82. W. Staels et al., Comprehensive alpha, beta, and delta cell transcriptomics reveal an association of cellular aging with MHC class I upregulation. Mol Metab 87, 101990 (2024).

83. T. Dos Santos et al., Altered immune and metabolic molecular pathways drive islet cell dysfunction in human type 1 diabetes. J Clin Invest, (2025).

84. X. Q. Dai et al., Heterogenous impairment of alpha cell function in type 2 diabetes is linked to cell maturation state. Cell Metabolism 34, 256-+ (2022).

85. V. D. Longo, R. M. Anderson, Nutrition, longevity and disease: From molecular mechanisms to interventions. Cell 185, 1455–1470 (2022).

86. J. A. Mattison et al., Impact of caloric restriction on health and survival in rhesus monkeys from the NIA study. Nature 489, 318–321 (2012).

87. A. E. Civitarese et al., Calorie restriction increases muscle mitochondrial biogenesis in healthy humans. PLoS Med 4, e76 (2007).

88. M. E. C. da Amaral et al., Caloric restriction recovers impaired beta-cell-beta-cell gap junction coupling, calcium oscillation coordination, and insulin secretion in prediabetic mice. American Journal of Physiology-Endocrinology and Metabolism 319, E709–E720 (2020).

89. C. J. Sheng et al., Reversibility of beta-Cell-Specific Transcript Factors Expression by Long-Term Caloric Restriction in db/db Mouse. Journal of Diabetes Research 2016, 11 (2016).

90. S. I. Tschen, S. Dhawan, T. Gurlo, A. Bhushan, Age-dependent decline in beta-cell proliferation restricts the capacity of beta-cell regeneration in mice. Diabetes 58, 1312–1320 (2009).

91. M. E. Ritchie et al., limma powers differential expression analyses for RNA-sequencing and microarray studies. Nucleic acids research 43, e47–e47 (2015).

92. M. I. Love, W. Huber, S. Anders, Moderated estimation of fold change and dispersion for RNA-seq data with DESeq2. Genome Biol 15, 550 (2014).

93. Y. Hao et al., Dictionary learning for integrative, multimodal and scalable single-cell analysis. Nat Biotechnol 42, 293–304 (2024).

94. Y. J. Wang et al., Single-Cell Mass Cytometry Analysis of the Human Endocrine Pancreas. Cell Metabolism 24, 616–626 (2016).

95. R. J. Colman et al., Caloric restriction delays disease onset and mortality in rhesus monkeys. Science 325, 201–204 (2009).

