## Supplementary Figure for "Alpha cell inflammation during human pancreas aging and type 2 diabetes and its reversal by calorie restriction in mice"

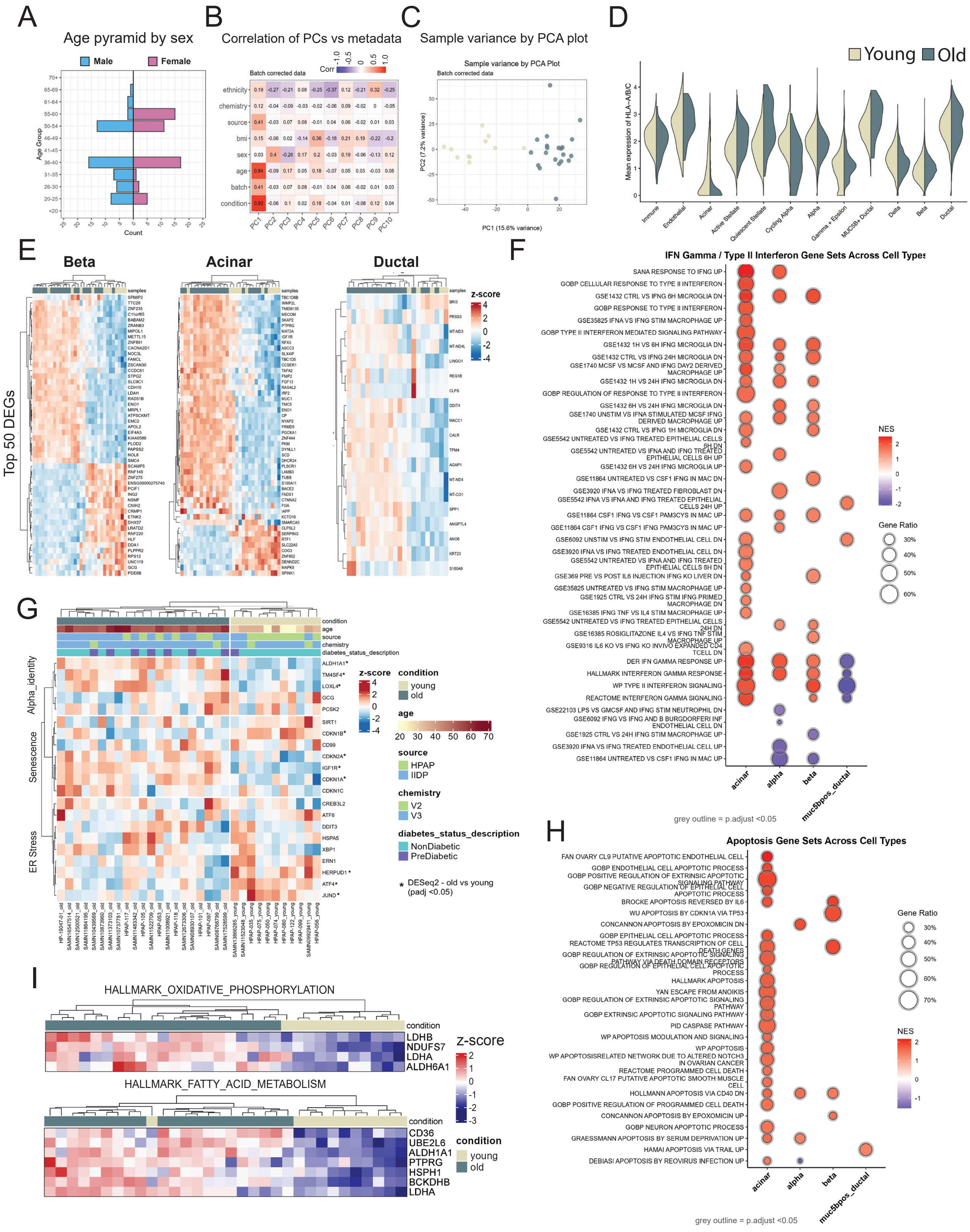

A
Age pyramid by sex
Male
Female
Age Group
Count
B
Correlation of PCs vs metadata
Batch corrected data
Corr
ethnicity
chemistry
source
bmi
sex
age
batch
condition
C
Sample variance by PCA plot
Sample variance by PCA Plot
Batch corrected data
PC1 (15.6% variance)
PC2 (7.2% variance)
D
Young
Old
Mean expression of HLA-A/B/C
Immune
Endothelial
Acinar
Active Stellate
Quiescent Stellate
Cycling Alpha
Alpha
Gamma + Epsilon
MUC5B+ Ductal
Delta
Beta
Ductal
E
Beta
Acinar
Ductal
Top 50 DEGs
z-score
F
IFN Gamma / Type II Interferon Gene Sets Across Cell Types
SANA RESPONSE TO IFNG UP
GOBP CELLULAR RESPONSE TO TYPE II INTERFERON
GSE1432 CTRL VS IFNG 6H MICROGLIA DN
GOBP RESPONSE TO TYPE II INTERFERON
GSE35825 IFNA VS IFNG STIM MACROPHAGE UP
GOBP TYPE II INTERFERON MEDIATED SIGNALING PATHWAY
GSE1432 1H VS 6H IFNG MICROGLIA DN
GSE1432 CTRL VS IFNG 24H MICROGLIA DN
GSE1740 MCSF VS MCSF AND IFNG DAY2 DERIVED MACROPHAGE UP
GSE1432 1H VS 24H IFNG MICROGLIA DN
GOBP REGULATION OF RESPONSE TO TYPE II INTERFERON
GSE1432 6H VS 24H IFNG MICROGLIA DN
GSE1740 UNSTIM VS IFNA STIMULATED MCSF IFNG DERIVED MACROPHAGE UP
GSE1432 CTRL VS IFNG 1H MICROGLIA DN
GSE5542 UNTREATED VS IFNG TREATED EPITHELIAL CELLS 6H DN
GSE5542 UNTREATED VS IFNA AND IFNG TREATED EPITHELIAL CELLS 6H UP
GSE1432 6H VS 24H IFNG MICROGLIA UP
GSE11864 UNTREATED VS CSF1 IFNG IN MAC DN
GSE3920 IFNA VS IFNG TREATED FIBROBLAST DN
GSE5542 IFNA VS IFNA AND IFNG TREATED EPITHELIAL CELLS 24H UP
GSE11864 CSF1 IFNG VS CSF1 PAM3CYS IN MAC UP
GSE11864 CSF1 IFNG VS CSF1 IFNG PAM3CYS IN MAC UP
GSE6092 UNSTIM VS IFNG STIM ENDOTHELIAL CELL DN
GSE3920 IFNA VS IFNG TREATED ENDOTHELIAL CELL DN
GSE5542 UNTREATED VS IFNA AND IFNG TREATED EPITHELIAL CELLS 6H DN
GSE369 PRE VS POST IL6 INJECTION IFNG KO LIVER DN
GSE35825 UNTREATED VS IFNG STIM MACROPHAGE UP
GSE1925 CTRL VS 24H IFNG STIM IFNG PRIMED MACROPHAGE DN
GSE16385 IFNG TNF VS IL4 STIM MACROPHAGE UP
GSE5542 UNTREATED VS IFNG TREATED EPITHELIAL CELLS 24H DN
GSE16385 ROSIGLITAZONE IL4 VS IFNG TNF STIM MACROPHAGE UP
GSE9316 IL6 KO VS IFNG KO INVIVO EXPANDED CD4 TCELL DN
DER IFN GAMMA RESPONSE UP
HALLMARK INTERFERON GAMMA RESPONSE
WP TYPE II INTERFERON SIGNALING
REACTOME INTERFERON GAMMA SIGNALING
GSE22103 LPS VS GMCSF AND IFNG STIM NEUTROPHIL DN
GSE6092 IFNG VS IFNG AND B BURGDORFERI INF ENDOTHELIAL CELL DN
GSE1925 CTRL VS 24H IFNG STIM MACROPHAGE UP
GSE3920 IFNA VS IFNG TREATED ENDOTHELIAL CELL UP
GSE11864 UNTREATED VS CSF1 IFNG IN MAC UP
acinar
alpha
beta
muc5bpos_ductal
NES
Gene Ratio
30%
40%
50%
60%
grey outline = p.adjust <0.05
G
condition
age
source
chemistry
diabetes_status_description
Alpha_identity
Senescence
ER Stress
z-score
condition
young
old
age
source
HPAP
IIDP
chemistry
V2
V3
diabetes_status_description
NonDiabetic
PreDiabetic
* DESeq2 - old vs young (padj <0.05)
H
Apoptosis Gene Sets Across Cell Types
FAN OVARY CL9 PUTATIVE APOPTOTIC ENDOTHELIAL CELL
GOBP ENDOTHELIAL CELL APOPTOTIC PROCESS
GOBP POSITIVE REGULATION OF EXTRINSIC APOPTOTIC SIGNALING PATHWAY
GOBP NEGATIVE REGULATION OF EPITHELIAL CELL APOPTOTIC PROCESS
BROCKE APOPTOSIS REVERSED BY IL6
WU APOPTOSIS BY CDKN1A VIA TP53
CONCANNON APOPTOSIS BY EPOXOMICIN DN
GOBP EPITHELIAL CELL APOPTOTIC PROCESS
REACTOME TP53 REGULATES TRANSCRIPTION OF CELL DEATH GENES
GOBP REGULATION OF EXTRINSIC APOPTOTIC SIGNALING PATHWAY VIA DEATH DOMAIN RECEPTORS
GOBP REGULATION OF EPITHELIAL CELL APOPTOTIC PROCESS
HALLMARK APOPTOSIS
YAN ESCAPE FROM ANOIKIS
GOBP REGULATION OF EXTRINSIC APOPTOTIC SIGNALING PATHWAY
GOBP EXTRINSIC APOPTOTIC SIGNALING PATHWAY
PID CASPASE PATHWAY
WP APOPTOSIS MODULATION AND SIGNALING
WP APOPTOSIS
WP APOPTOSISRELATED NETWORK DUE TO ALTERED NOTCH3 IN OVARIAN CANCER
REACTOME PROGRAMMED CELL DEATH
FAN OVARY CL17 PUTATIVE APOPTOTIC SMOOTH MUSCLE CELL
HOLLMANN APOPTOSIS VIA CD40 DN
GOBP POSITIVE REGULATION OF PROGRAMMED CELL DEATH
CONCANNON APOPTOSIS BY EPOXOMICIN UP
GOBP NEURON APOPTOTIC PROCESS
GRAESSMANN APOPTOSIS BY SERUM DEPRIVATION UP
HAMAI APOPTOSIS VIA TRAIL UP
DEBIASI APOPTOSIS BY REOVIRUS INFECTION UP
acinar
alpha
beta
muc5bpos_ductal
Gene Ratio
NES
grey outline = p.adjust <0.05
I
HALLMARK_OXIDATIVE_PHOSPHORYLATION
condition
LDHB
NDUFS7
LDHA
ALDH6A1
z-score
HALLMARK_FATTY_ACID_METABOLISM
condition
CD36
UBE2L6
ALDH1A1
PTPRG
HSPH1
BCKDHB
LDHA
condition
young
old

**Supplementary Figure S1. (A)** Age and sex stratification for young (n=11; F=6, M=5) and old (n=21; F=11, M=10) ND donors for scRNA-seq integration, analysis, and SCENIC TF activity. **(B)** Batch correction improves sample clustering by experimental condition. Principal component analysis (PCA) of pseudobulk RNAseq data from pancreatic alpha cells across 32 donors. **(C)** Following RUVSeq batch correction PCA, technical correlations are substantially reduced while the age group condition correlation remains high at r=0.92, demonstrating effective removal of unwanted technical variation while preserving the biological signal of interest. Correlation heatmaps display Pearson correlation coefficients between the first 10 principal components and sample metadata variables. **(D)** Violin plots displaying HLA-ABC gene expression levels in pancreatic cells of young and old adult islets. **(E)** Heatmap displaying the differential gene expression of beta cells, acinar cells, and ductal cells in young versus old donors. **(F)** GSEA enrichment dotplot heatmap demonstrating the level of activation of several IFN-γ related pathways in alpha cells, beta cells, acinar cells, and ductal cells. **(G)** Hierarchical clustering heatmap displaying the relative expression levels of hallmark genes for alpha cell identity, senescence, and ER stress pathways. * denotes DESeq2 adjusted p values < 0.05 between old vs young groups. **(H)** GSEA enrichment dotplot heatmap demonstrating the level of activation of several apoptosis-related pathways in alpha cells, beta cells, acinar cells, and ductal cells. **(I)** Heatmap displaying the differentially regulated gene expression associated with oxidative phosphorylation and fatty acid metabolism in alpha cells from young versus old donors.

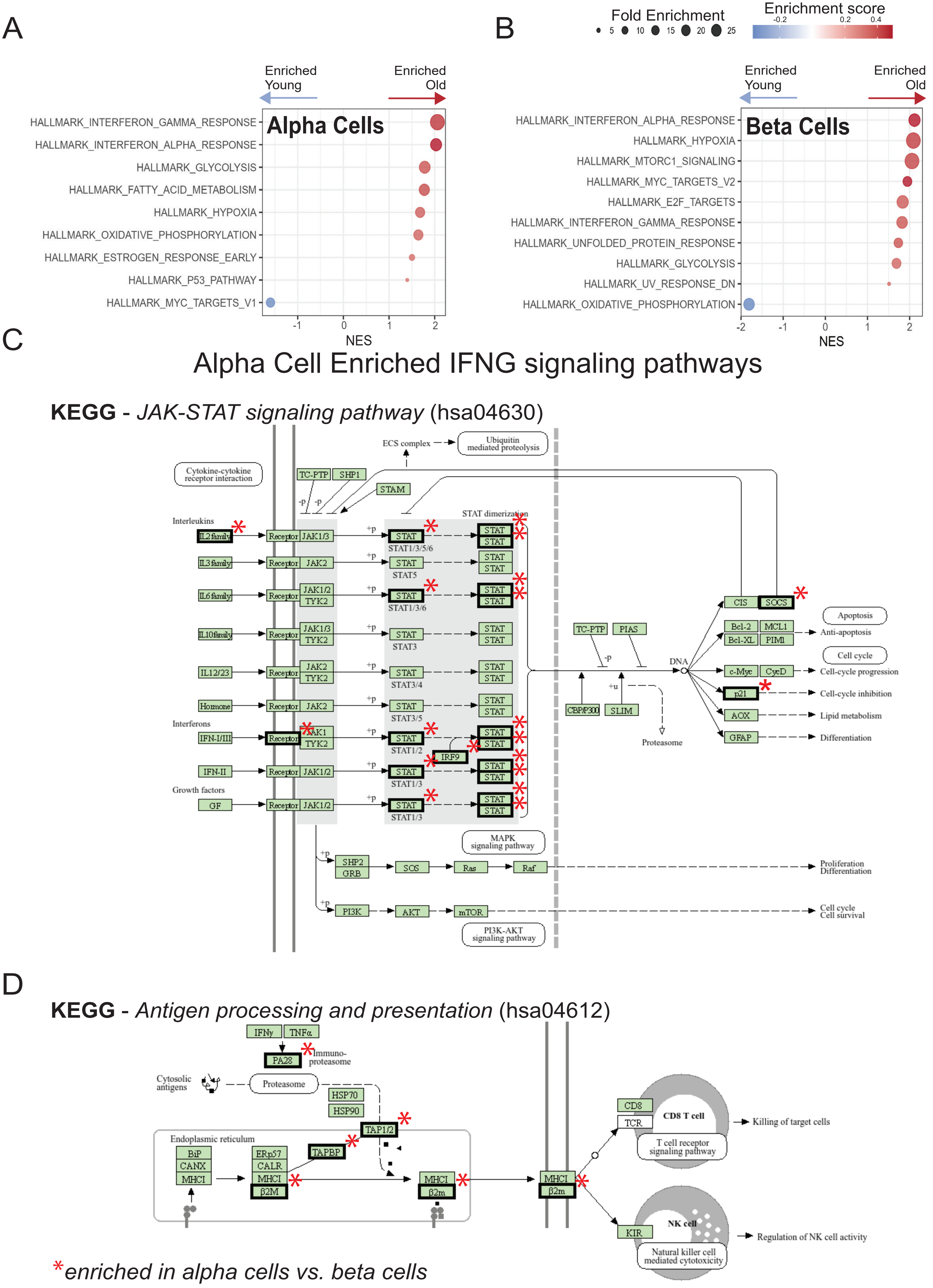

A
B
Fold Enrichment
5 10 15 20 25
Enrichment score
-0.2 0.2 0.4
Enriched Young
Enriched Old
Alpha Cells
HALLMARK_INTERFERON_GAMMA_RESPONSE
HALLMARK_INTERFERON_ALPHA_RESPONSE
HALLMARK_GLYCOLYSIS
HALLMARK_FATTY_ACID_METABOLISM
HALLMARK_HYPOXIA
HALLMARK_OXIDATIVE_PHOSPHORYLATION
HALLMARK_ESTROGEN_RESPONSE_EARLY
HALLMARK_P53_PATHWAY
HALLMARK_MYC_TARGETS_V1
-1 0 1 2
NES
Beta Cells
HALLMARK_INTERFERON_ALPHA_RESPONSE
HALLMARK_HYPOXIA
HALLMARK_MTORC1_SIGNALING
HALLMARK_MYC_TARGETS_V2
HALLMARK_E2F_TARGETS
HALLMARK_INTERFERON_GAMMA_RESPONSE
HALLMARK_UNFOLDED_PROTEIN_RESPONSE
HALLMARK_GLYCOLYSIS
HALLMARK_UV_RESPONSE_DN
HALLMARK_OXIDATIVE_PHOSPHORYLATION
-2 -1 0 1 2
NES
C
Alpha Cell Enriched IFNG signaling pathways
KEGG - JAK-STAT signaling pathway (hsa04630)
ECS complex
Ubiquitin mediated proteolysis
Cytokine-cytokine receptor interaction
TC-PTP
SHP1
STAM
STAT dimerization
Interleukins
IL2 family
IL3 family
IL6 family
IL10 family
IL12/23
Hormone
Interferons
IFN-I/III
IFN-II
Growth factors
GF
Receptor
JAK1/3
JAK2
JAK1/2
TYK2
JAK1
STAT
STAT1/3/5/6
STAT5
STAT1/3/6
STAT3
STAT3/4
STAT3/5
STAT1/2
STAT1/3
IRF9
TC-PTP
PIAS
CBP/P300
SLIM
Proteasome
DNA
CIS
SOCS
Bcl-2
MCL1
Bcl-XL
PIM1
c-Myc
CycD
p21
AOX
GFAP
Apoptosis
Anti-apoptosis
Cell cycle
Cell-cycle progression
Cell-cycle inhibition
Lipid metabolism
Differentiation
MAPK signaling pathway
SHP2
GRB
SOS
Ras
Raf
Proliferation Differentiation
PI3K
AKT
mTOR
Cell cycle Cell survival
PI3K-AKT signaling pathway
D
KEGG - Antigen processing and presentation (hsa04612)
IFNγ
TNFα
PA28
Immuno-proteasome
Cytosolic antigens
Proteasome
HSP70
HSP90
TAP1/2
TAPBP
Endoplasmic reticulum
BiP
CANX
MHCI
ERp57
CALR
β2M
β2m
CD8
TCR
CD8 T cell
T cell receptor signaling pathway
Killing of target cells
KIR
NK cell
Natural killer cell mediated cytotoxicity
Regulation of NK cell activity
*enriched in alpha cells vs. beta cells

**Supplementary Figure S2. (A-B)** Pathway enrichment scores (NES) after GSEA analysis of genes differentially regulate by age in alpha cells and beta cells, respectively. **(C-D)** Annotated KEGG pathway map highlighting the genes upregulated in old alpha cells and that belong to the JAK-STAT signaling pathway and antigen processing and presentation, respectively. Black bounding boxes and asterisk mark upregulated genes and enriched in alpha cells versus beta cells.

### HPAP - Vanderbilt

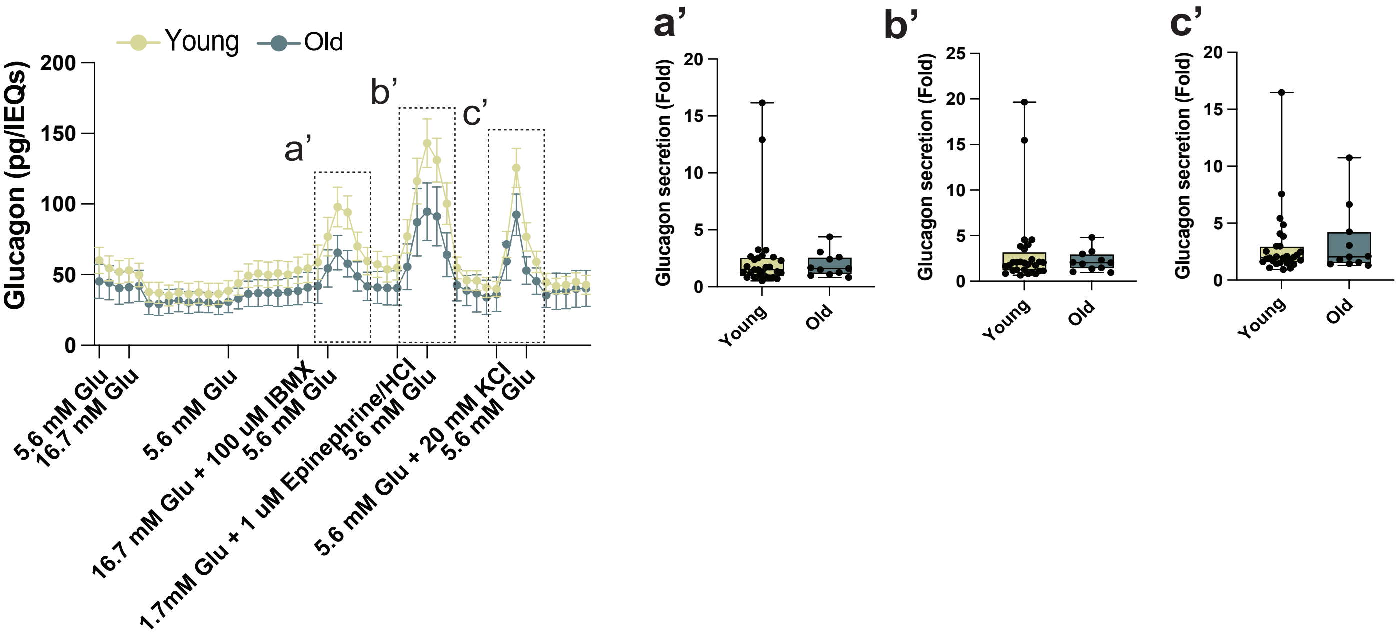

**Supplementary Figure S3.** Isolated human islet perifusion assay including stimulation with glutamine, glucose, epinephrine, and KCL. Boxplots in Panel D, a'-c' are mean ± IQR with minimum and maximum values.

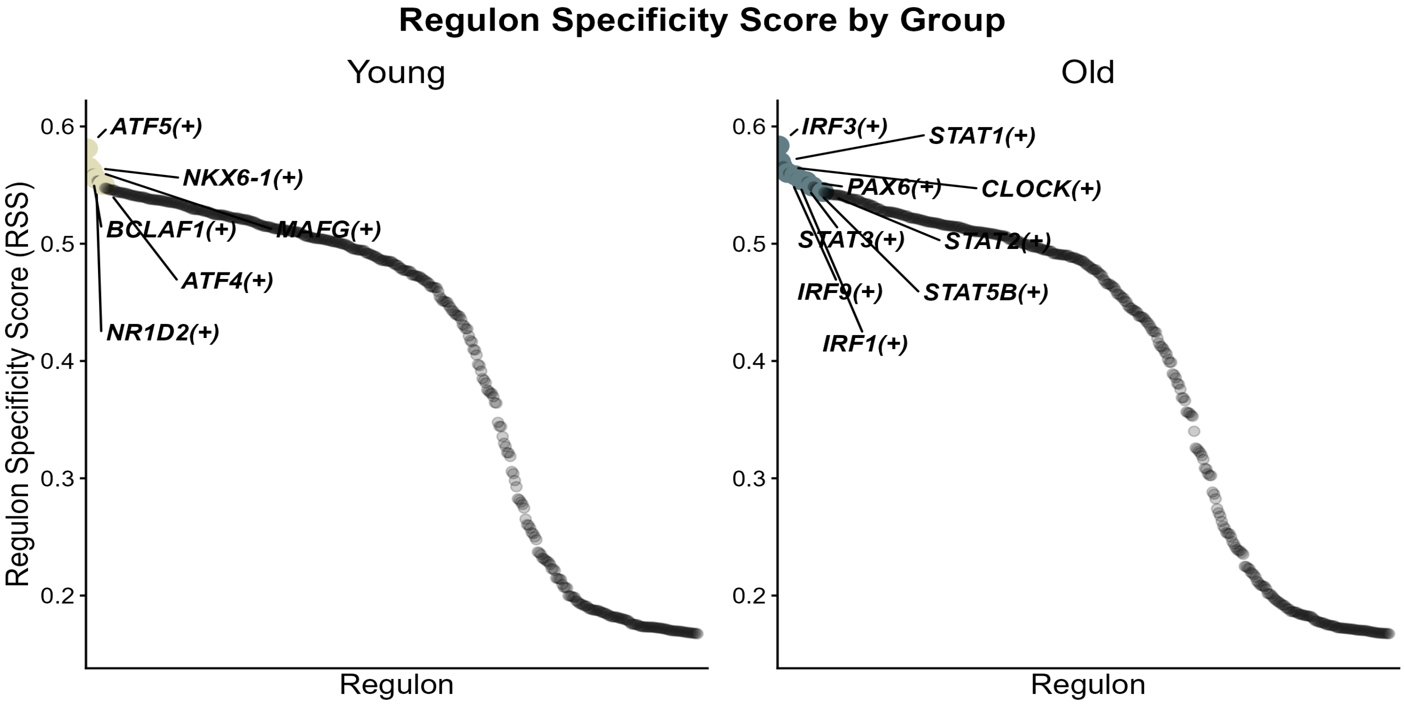
Regulon Specificity Score by Group
Young
Old
Regulon Specificity Score (RSS)
Regulon
0.6
0.5
0.4
0.3
0.2
ATF5(+)
NKX6-1(+)
BCLAF1(+)
MAFG(+)
ATF4(+)
NR1D2(+)
IRF3(+)
STAT1(+)
PAX6(+)
CLOCK(+)
STAT3(+)
STAT2(+)
IRF9(+)
STAT5B(+)
IRF1(+)

**Supplementary Figure S4.** Regulon Specificity Score (RSS) in young (left) and old (right) adult alpha cells. Highlighted and labeled features represent transcription factors within the top 10% of RSS scores in each group.

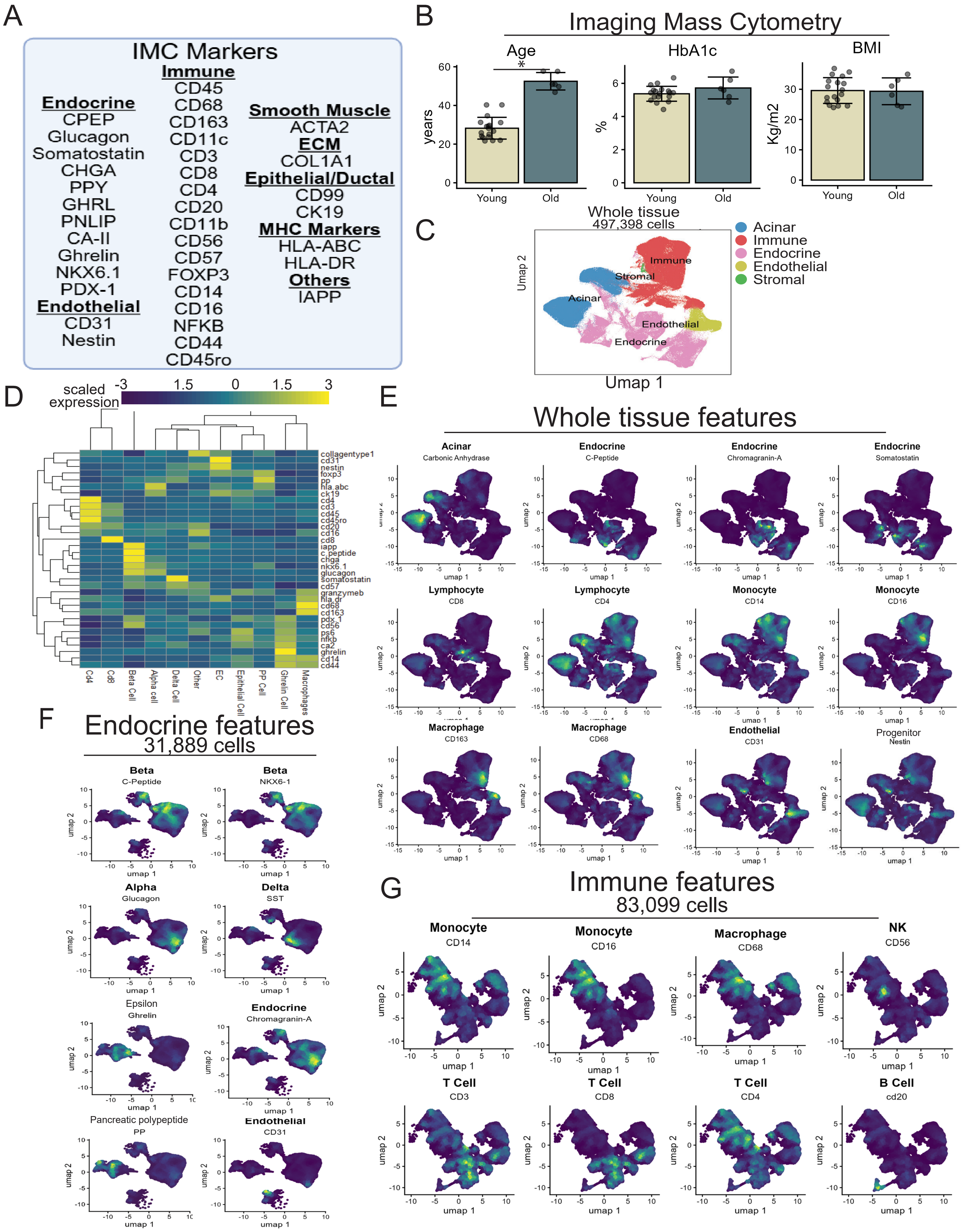
A
IMC Markers
Endocrine
CPEP
Glucagon
Somatostatin
CHGA
PPY
GHRL
PNLIP
CA-II
Ghrelin
NKX6.1
PDX-1
Endothelial
CD31
Nestin
Immune
CD45
CD68
CD163
CD11c
CD3
CD8
CD4
CD20
CD11b
CD56
CD57
FOXP3
CD14
CD16
NFKB
CD44
CD45ro
Smooth Muscle
ACTA2
ECM
COL1A1
Epithelial/Ductal
CD99
CK19
MHC Markers
HLA-ABC
HLA-DR
Others
IAPP
B
Imaging Mass Cytometry
Age
HbA1c
BMI
years
%
Kg/m2
Young
Old
*
C
Whole tissue
497,398 cells
Acinar
Immune
Endocrine
Endothelial
Stromal
Umap 1
Umap 2
D
scaled expression
-3
1.5
0
1.5
3
collagentype1
cd31
nestin
foxp3
pp
hla.abc
ck19
cd4
cd3
cd45
cd45ro
cd20
cd16
cd8
iapp
c.peptide
chga
nkx6.1
glucagon
somatostatin
cd57
granzymeb
hla.dr
cd68
cd163
pdx.1
cd56
ps6
nfkb
ca2
ghrelin
cd14
cd44
Cd4
Cd8
Beta Cell
Alpha cell
Delta Cell
Other
EC
Epithelial Cell
PP Cell
Ghrelin Cell
Macrophages
E
Whole tissue features
Acinar
Carbonic Anhydrase
Endocrine
C-Peptide
Endocrine
Chromagranin-A
Endocrine
Somatostatin
Lymphocyte
CD8
Lymphocyte
CD4
Monocyte
CD14
Monocyte
CD16
Macrophage
CD163
Macrophage
CD68
Endothelial
CD31
Progenitor
Nestin
umap 1
umap 2
F
Endocrine features
31,889 cells
Beta
C-Peptide
Beta
NKX6-1
Alpha
Glucagon
Delta
SST
Epsilon
Ghrelin
Endocrine
Chromagranin-A
Pancreatic polypeptide
PP
Endothelial
CD31
G
Immune features
83,099 cells
Monocyte
CD14
Monocyte
CD16
Macrophage
CD68
NK
CD56
T Cell
CD3
T Cell
CD8
T Cell
CD4
B Cell
cd20

**Supplementary Figure S5. (A)** Imaging Mass Cytometry (IMC) antibody and marker list. **(B)** Donors were discordant for age but presented similar HbA1c and BMI. **(C)** UMAP subsets for acinar, immune, endothelial, endocrine, and stromal cell populations. **(D)** Clustering of all major cell types presented using mean z-normalized marker expression for each cell-type. **(E)** Feature plots for acinar (carbonic anhydrase [CA2]), endocrine (chromogranin-A, somatostatin, C-peptide), immune (CD4, CD8, CD14, CD16, CD68, CD163), and endothelial (CD31), and progenitor (nestin) markers. **(F)** Endocrine cell features for beta cells (C-Peptide, NKX6-1, alpha cells (glucagon), delta cells (somatostatin), epsilon cells (ghrelin), chromogranin-A, pancreatic polypeptide (PP), and endothelial cells (CD31). **(G)** Immune cell features for monocytes (CD14, CD16), macrophages (CD68, NK cells (CD56), T cells (CD3, CD8, CD4) and B cells (CD20). Data are mean±SEM in panel B.

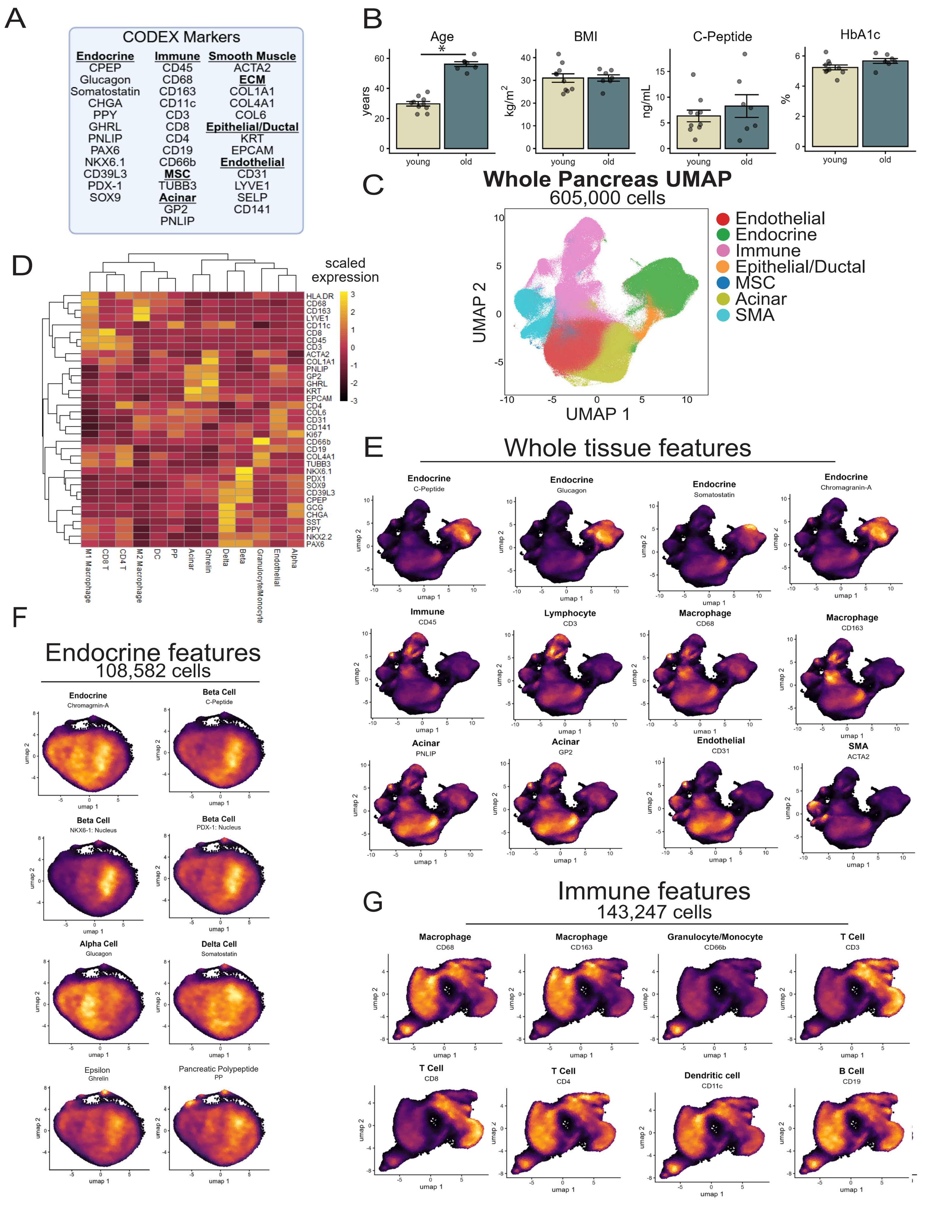
A
CODEX Markers
Endocrine
CPEP
Glucagon
Somatostatin
CHGA
PPY
GHRL
PNLIP
PAX6
NKX6.1
CD39L3
PDX-1
SOX9
Immune
CD45
CD68
CD163
CD11c
CD3
CD8
CD4
CD19
CD66b
MSC
TUBB3
Acinar
GP2
PNLIP
Smooth Muscle
ACTA2
ECM
COL1A1
COL4A1
COL6
Epithelial/Ductal
KRT
EPCAM
Endothelial
CD31
LYVE1
SELP
CD141
B
Age
BMI
C-Peptide
HbA1c
years
kg/m2
ng/mL
%
young
old
C
Whole Pancreas UMAP
605,000 cells
UMAP 1
UMAP 2
Endothelial
Endocrine
Immune
Epithelial/Ductal
MSC
Acinar
SMA
D
scaled expression
HLA.DR
CD68
CD163
LYVE1
CD11c
CD8
CD45
CD3
ACTA2
COL1A1
PNLIP
GP2
GHRL
KRT
EPCAM
CD4
COL6
CD31
CD141
Ki67
CD66b
CD19
COL4A1
TUBB3
NKX6.1
PDX1
SOX9
CD39L3
CPEP
GCG
CHGA
SST
PPY
NKX2.2
PAX6
M1 Macrophage
CD8 T
CD4 T
M2 Macrophage
DC
PP
Acinar
Ghrelin
Delta
Beta
Granulocyte/Monocyte
Endothelial
Alpha
E
Whole tissue features
Endocrine
C-Peptide
Endocrine
Glucagon
Endocrine
Somatostatin
Endocrine
Chromagranin-A
Immune
CD45
Lymphocyte
CD3
Macrophage
CD68
Macrophage
CD163
Acinar
PNLIP
Acinar
GP2
Endothelial
CD31
SMA
ACTA2
F
Endocrine features
108,582 cells
Endocrine
Chromagnin-A
Beta Cell
C-Peptide
Beta Cell
NKX6-1: Nucleus
Beta Cell
PDX-1: Nucleus
Alpha Cell
Glucagon
Delta Cell
Somatostatin
Epsilon
Ghrelin
Pancreatic Polypeptide
PP
G
Immune features
143,247 cells
Macrophage
CD68
Macrophage
CD163
Granulocyte/Monocyte
CD66b
T Cell
CD3
T Cell
CD8
T Cell
CD4
Dendritic cell
CD11c
B Cell
CD19
umap 1
umap 2

**Supplementary Figure S6. (A)** CODEX antibody and marker list. **(B)** Donors were discordant for age, but presented similar BMI, circulating C-peptide, and HbA1c. **(C)** UMAP subsets for endothelial, endocrine, immune, epithelial/ductal, mesenchymal stem cell (MSC), acinar, and smooth muscle actin (SMA) cell populations. **(D)** Clustering of all major cell types presented using mean z-normalized marker expression for each cell-type. **(E)** Feature plots for endocrine (C-peptide, glucagon, somatostatin, chromogranin-A), pan-immune (CD45), lymphocytes (CD3), macrophages (CD68, CD163), acinar (pancreatic lipase [PNLIP], glycoprotein 2 [GP2]), endothelial (CD31), and SMA (ACTA2) cell populations. **(F)** Endocrine cell features for pan-neuroendocrine cells (chromogranin-A), beta cells (C-peptide, NKX6-1, and PDX-1), alpha cells (glucagon), delta cells (somatostatin), epsilon (ghrelin), and pancreatic polypeptide cells (PP). **(G)** Immune cell features for macrophages (CD68, CD163), granulocytes/monocytes (CD66b), T cells (CD3, CD8, CD4), dendritic cells (CD11c), and B cells (CD19). Data are mean±SEM in panel B.

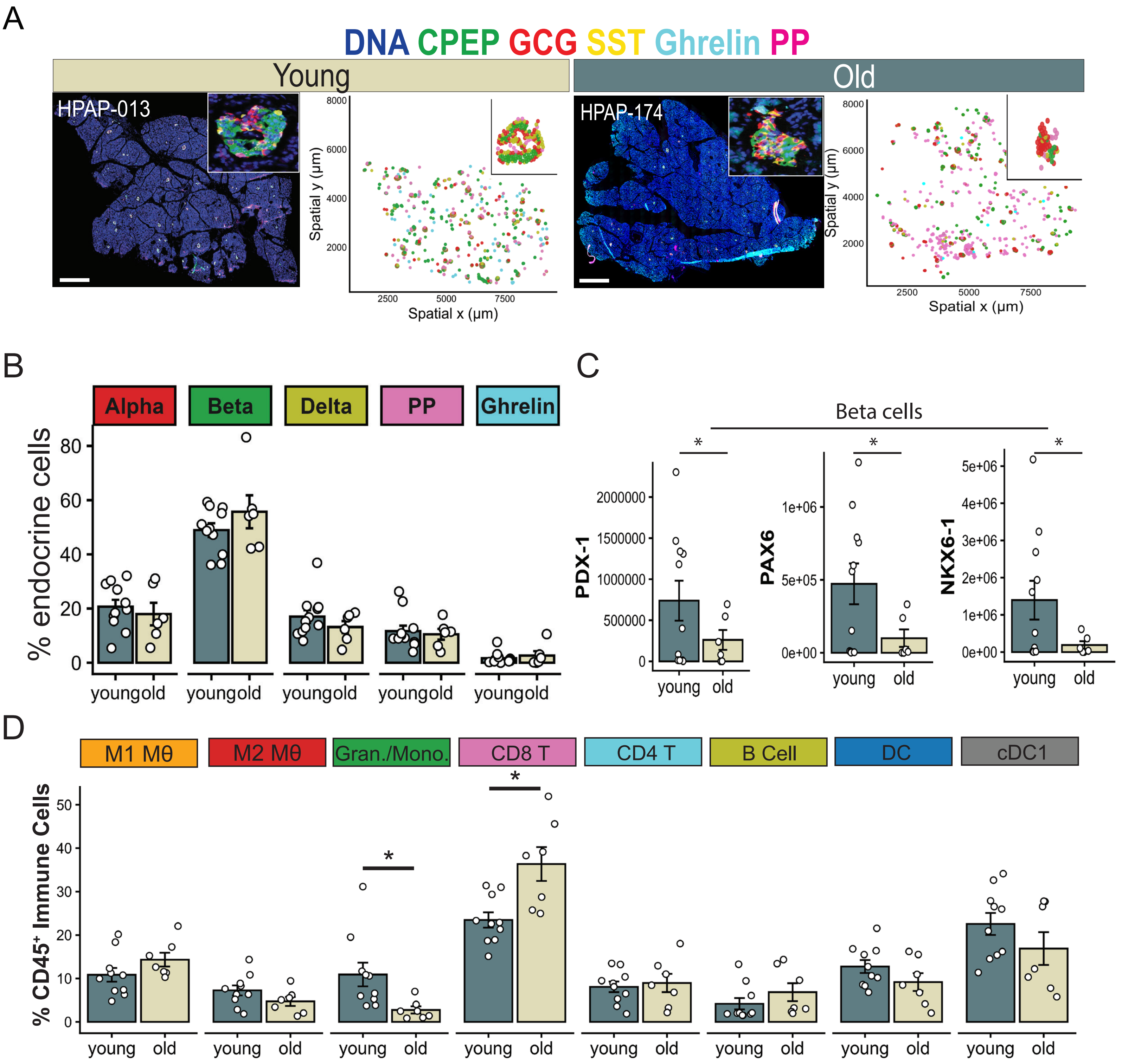

A
DNA CPEP GCG SST Ghrelin PP
Young
Old
HPAP-013
HPAP-174
Spatial y (μm)
Spatial x (μm)
B
Alpha
Beta
Delta
PP
Ghrelin
% endocrine cells
young old
C
Beta cells
PDX-1
PAX6
NKX6-1
young
old
D
M1 Mθ
M2 Mθ
Gran./Mono.
CD8 T
CD4 T
B Cell
DC
cDC1
% CD45⁺ Immune Cells

**Supplementary Figure S7. (A)** Representative CODEX images and spatial reconstructions for islet endocrine subsets. Scale bar = 2000µm. **(B)** Relative endocrine cell proportions for alpha cells ($GCG^+$), beta cells (C-peptide$^+$), delta cells ($SST^+$), Pancreatic polypeptide ($PP^+$), and epsilon cells (ghrelin$^+$). **(C)** Paired expression for beta cell identity markers, PDX-1, PAX6, and NKX6-1. **(D)** Relative immune cell populations for M1 macrophages ($CD68^+CD163^-$; Mθ), M2 macrophages ($CD68^+CD163^+$), granulocytes/monocytes ($CD66b^+$), CD8 T cells ($CD3^+CD8^+$), CD4 T cells ($CD3^+CD4^+$), B cells ($CD19^+$, $CD20^+$), dendritic cells (DC: $CD11c^+$), and conventional DC (cDCs; $CD141^+$). n=8-11/group. Welch's two-sample t-test compared young vs. old donor groups. Data are mean±SEM. *$p<0.05$, young vs old ND donors.

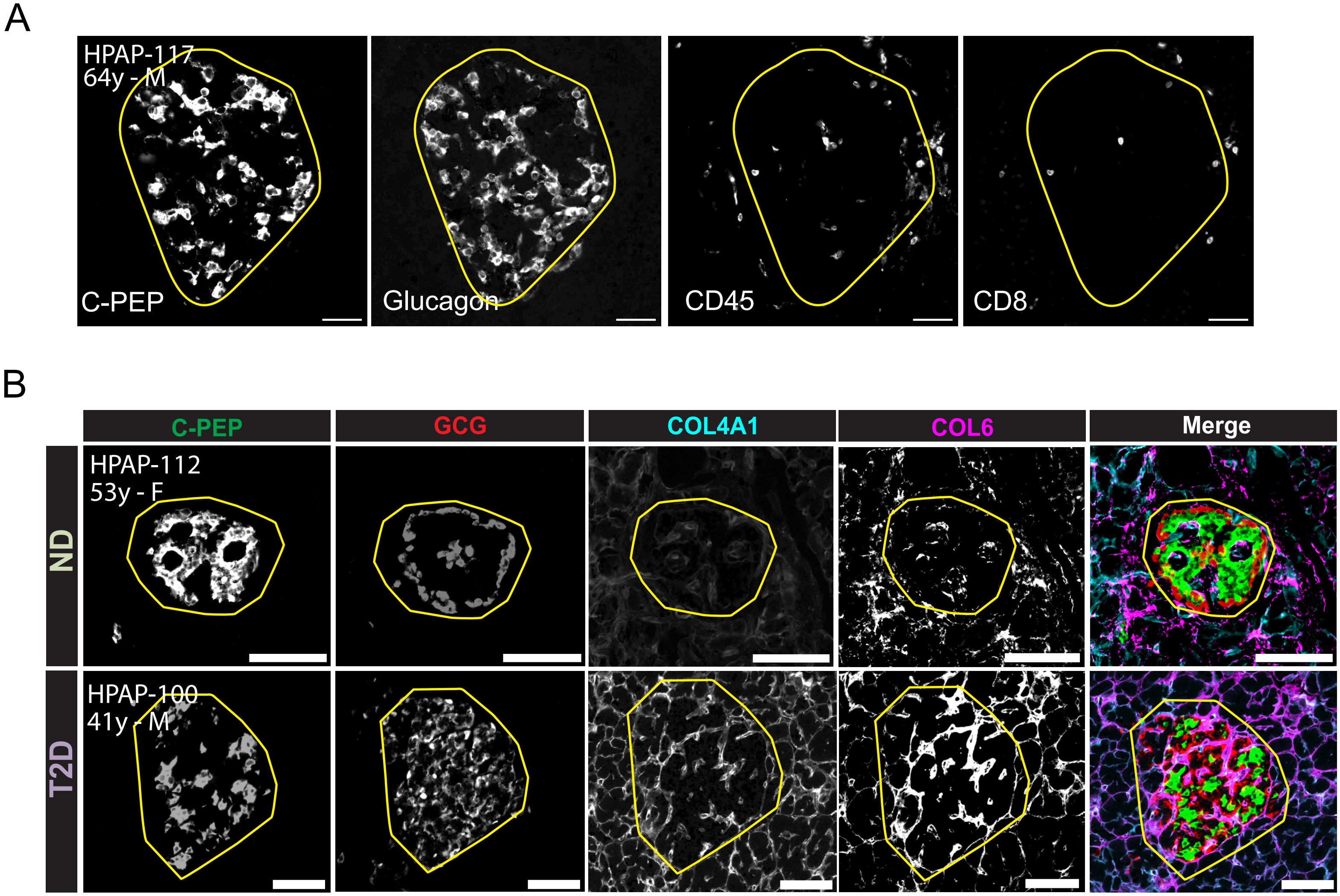

A
HPAP-117
64y - M
C-PEP
Glucagon
CD45
CD8
B
C-PEP
GCG
COL4A1
COL6
Merge
ND
HPAP-112
53y - F
T2D
HPAP-100
41y - M

**Supplementary Figure S8. (A)** Representative images of islets (C-PEP, glucagon) and $CD45^{+}CD8^{+}$ T cells, corresponding to Figure 4E. Representative islet showing beta cells (C-PEP, green), alpha cells (GCG, red), and collagens (COL4A1, cyan; COL6, magenta). Scale bar = 100μm, corresponding to Figure 5C.

### CD8 T Cell Phenotyping

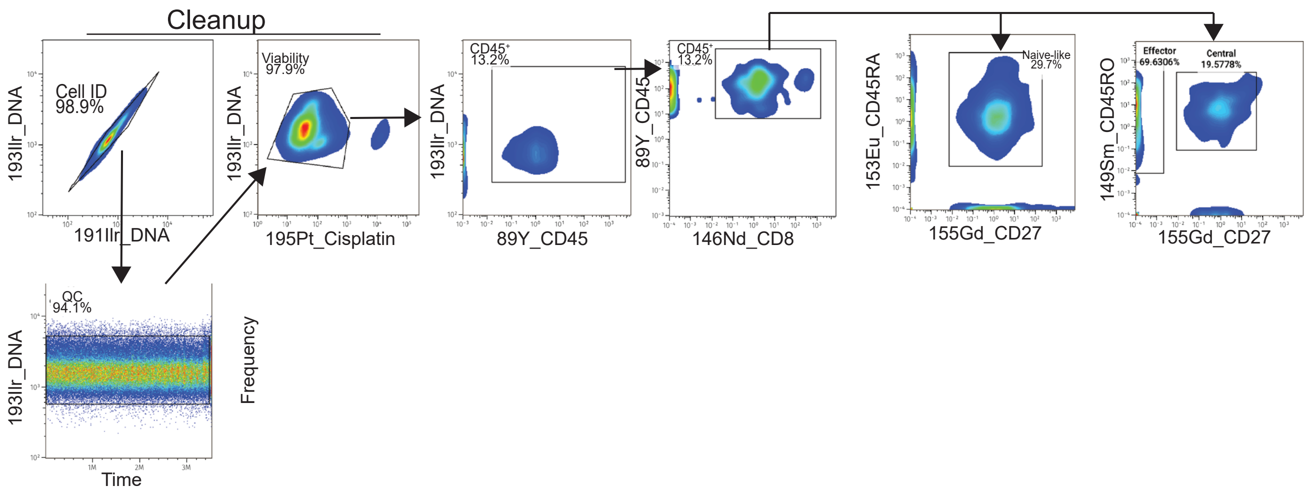

**Supplementary Figure S9.** Representative Cytometry Time of Flight (CyToF) gating strategy for $CD45^+CD8^+$ T cells, including naïve ($CD27^+CD45RA^+$), memory ($CD27^+CD45RA^-$), and terminal effector memory ($CD27^-CD45RA^+$) phenotypes.

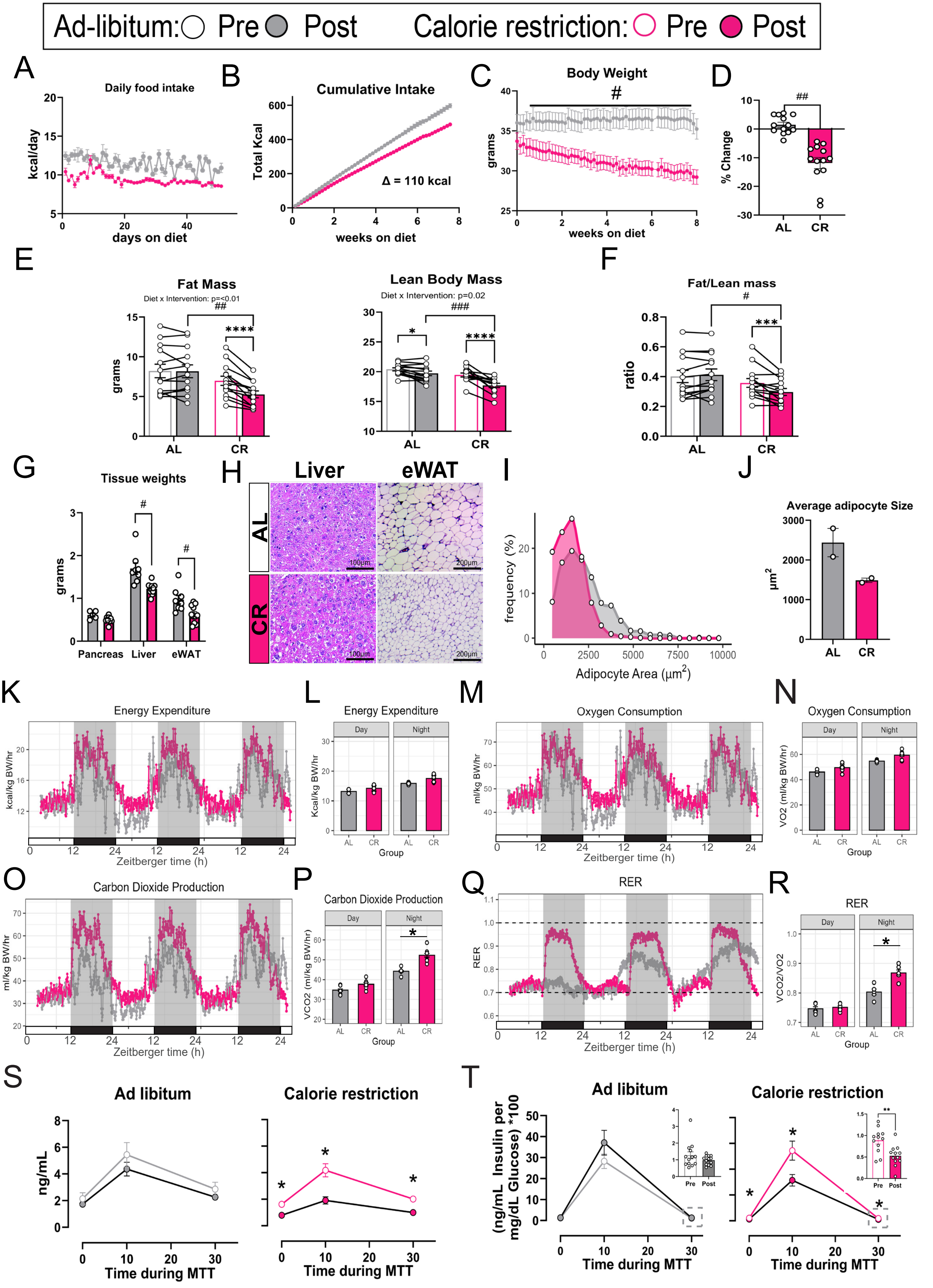
Ad-libitum: Pre Post Calorie restriction: Pre Post
A
Daily food intake
kcal/day
days on diet
B
Cumulative Intake
Total Kcal
Δ = 110 kcal
weeks on diet
C
Body Weight
#
grams
weeks on diet
D
##
% Change
AL
CR
E
Fat Mass
Diet x Intervention: p=<0.01
##
****
grams
Lean Body Mass
Diet x Intervention: p=0.02
###
*
****
F
Fat/Lean mass
#
***
ratio
G
Tissue weights
#
#
grams
Pancreas
Liver
eWAT
H
Liver
eWAT
AL
CR
100μm
200μm
I
frequency (%)
Adipocyte Area (μm²)
J
Average adipocyte Size
μm²
K
Energy Expenditure
kcal/kg BW/hr
Zeitberger time (h)
L
Energy Expenditure
Day
Night
Kcal/kg BW/hr
Group
M
Oxygen Consumption
ml/kg BW/hr
Zeitberger time (h)
N
Oxygen Consumption
VO2 (ml/kg BW/hr)
Group
O
Carbon Dioxide Production
ml/kg BW/hr
Zeitberger time (h)
P
Carbon Dioxide Production
*
VCO2 (ml/kg BW/hr)
Group
Q
RER
Zeitberger time (h)
R
RER
*
VCO2/VO2
Group
S
Ad libitum
Calorie restriction
ng/mL
Time during MTT
T
Ad libitum
Calorie restriction
(ng/mL Insulin per mg/dL Glucose) *100
Time during MTT
Pre
Post
**

**Supplementary Figure S10. (A).** Energy intake, kcals, consumed daily. **(B)** cumulative energy intake over the intervention period. **(C)** Absolute measures of body weight over the intervention period. **(D)** Relative body weight change (% change from baseline) after 2 months CR. **(E)** Fat mass and lean body mass before and intervention. **(F)** Fat-to-lean mass ratio. **(G)** Tissue weights (pancreas, liver, eWAT), presented in grams. **(H)** Representative H&E image from liver and eWAT. Scale bar = 100µm. **(I)** Frequency of adipocytes from 250-10,000µm, bins=20. **(J)** Average adipocyte size. **(K)** Energy expenditure (kcals/kg body weight/hour) measured relative to total body weight collected every five minutes. **(L)** Energy expenditure aggregated into the day (light; 6:00am-6:00pm) and night (dark; 6:00pm-6:00am) phase. **(M)** Oxygen consumption (VO2) measured continuously relative to total body weight. **(N)** VO2 relative to body weight aggregated into day and night periods. **(O)** Carbon dioxide production (VCO2) measured continuously relative to total body weight. **(P)** VCO2 relative to body weight aggregated into day and night periods. **(Q)** Respiratory exchange ratio (RER; VCO2/VO2) measured continuously. **(R)** RER measures aggregated into day and night periods. **(S)** Insulin concentration during the MTT. **(T)** Insulin-to-glucose ratio during the MTT. Two-way ANOVA with main effects for diet (CR vs. AL) and time (day vs night) was completed with Tukey adjustments in panels J,L,N,P. n=11/group for panels A-D; n=9/group for panel E; n=2/group for panels F-H; n=5/group for panels I-P. Data are mean±SEM. * $p<0.05$, AL vs CR.

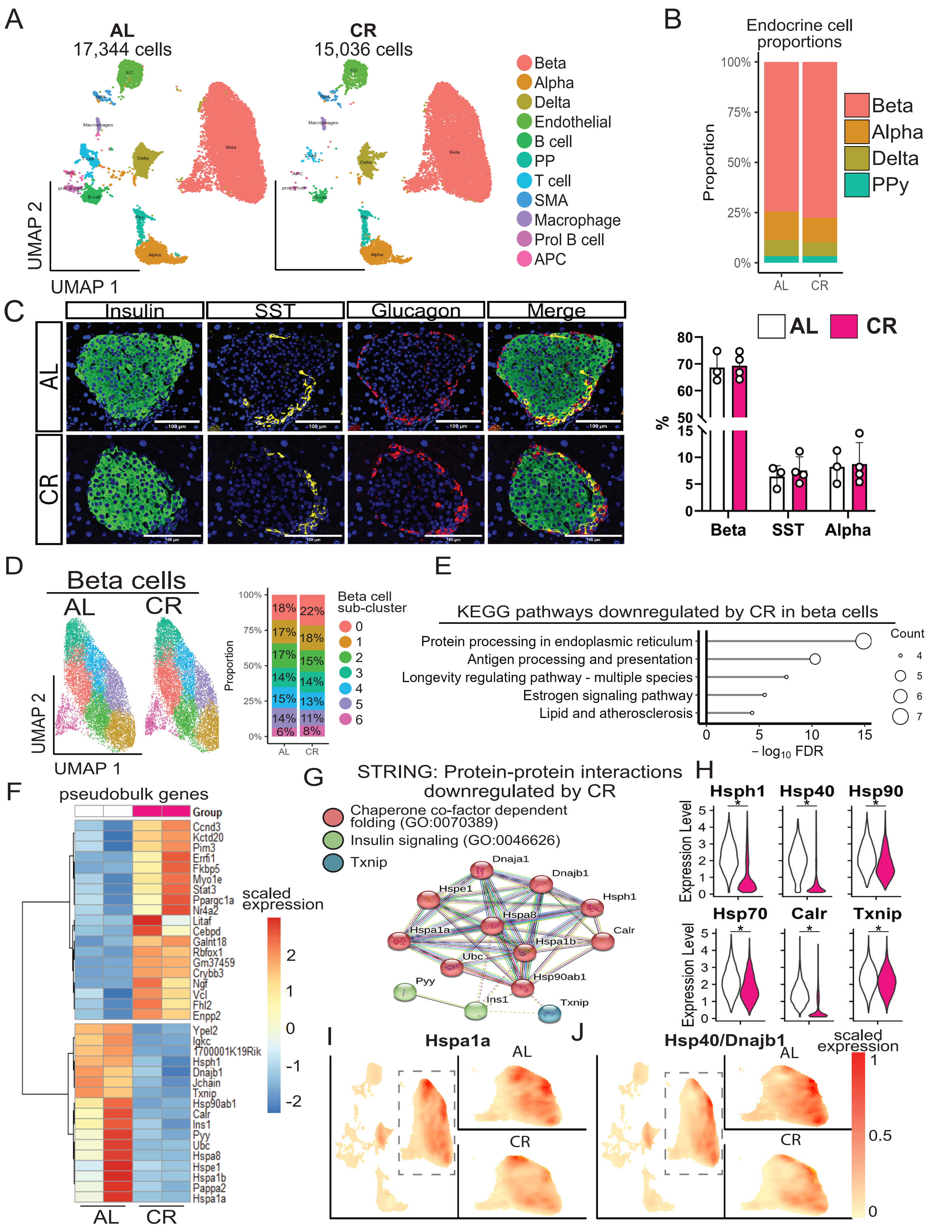

A
AL
17,344 cells
CR
15,036 cells
EC
SMA
Macrophages
T-cell
APC
prol_B-cell
B-cell
Delta
Beta
Ppy
Alpha
UMAP 2
UMAP 1
Beta
Alpha
Delta
Endothelial
B cell
PP
T cell
SMA
Macrophage
Prol B cell
APC
B
Endocrine cell proportions
Proportion
100%
75%
50%
25%
0%
AL
CR
Beta
Alpha
Delta
PPy
C
Insulin
SST
Glucagon
Merge
AL
CR
100 µm
AL
CR
%
80
70
60
50
15
10
5
0
Beta
SST
Alpha
D
Beta cells
AL
CR
UMAP 2
UMAP 1
Proportion
100%
75%
50%
25%
0%
18%
22%
17%
18%
17%
15%
14%
14%
15%
13%
14%
11%
6%
8%
AL
CR
Beta cell sub-cluster
0
1
2
3
4
5
6
E
KEGG pathways downregulated by CR in beta cells
Protein processing in endoplasmic reticulum
Antigen processing and presentation
Longevity regulating pathway - multiple species
Estrogen signaling pathway
Lipid and atherosclerosis
0
5
10
15
− log10 FDR
Count
4
5
6
7
F
pseudobulk genes
Group
Ccnd3
Kctd20
Pim3
Errfi1
Fkbp5
Myo1e
Stat3
Ppargc1a
Nr4a2
Litaf
Cebpd
Galnt18
Rbfox1
Gm37459
Crybb3
Ngf
Vcl
Fhl2
Enpp2
Ypel2
Igkc
1700001K19Rik
Hsph1
Dnajb1
Jchain
Txnip
Hsp90ab1
Calr
Ins1
Pyy
Ubc
Hspa8
Hspe1
Hspa1b
Pappa2
Hspa1a
AL
CR
scaled expression
2
1
0
-1
-2
G
STRING: Protein-protein interactions downregulated by CR
Chaperone co-factor dependent folding (GO:0070389)
Insulin signaling (GO:0046626)
Txnip
Dnaja1
Dnajb1
Hspe1
Hsph1
Hspa8
Hspa1a
Calr
Hspa1b
Ubc
Hsp90ab1
Pyy
Ins1
Txnip
H
Hsph1
Hsp40
Hsp90
Expression Level
5
4
3
2
1
0
*
Hsp70
Calr
Txnip
Expression Level
I
Hspa1a
AL
CR
J
Hsp40/Dnajb1
AL
CR
scaled expression
1
0.5
0

**Supplementary Figure S11. (A)** UMAP of 32,380 islet nuclei for single cell experiments. **(B)** Proportion of endocrine islet cells per group. **(C)** Representative islets illustrating beta cells (insulin), delta cells (somatostatin), and alpha cells (glucagon), and percent composition per islet. n=3-4 mice/group from 9-12 islets per mouse. Scale bar = 100µm. **(D)** Beta cell subclusters and the proportion within each subcluster. **(E)** KEGG pathways significantly downregulated by CR in beta cells. **(F)** Differential gene expression in beta cells by pseudobulk analysis. **(G)** Protein-protein interaction networks shows predicted and known associations among genes downregulated by CR (STRING). **(H)** Expression of beta cell genes related to protein processing and antigen presentation. **(I-J)** Density of Hspa1a (I) and Hsp40/Dnajb1 (J) for sequenced nuclei. KEGG, Kyoto encyclopedia of genes and genomes; UMAP, Uniform Manifold Approximation and Projection. *$p < 0.05$ for dietary intervention (AL vs. CR).

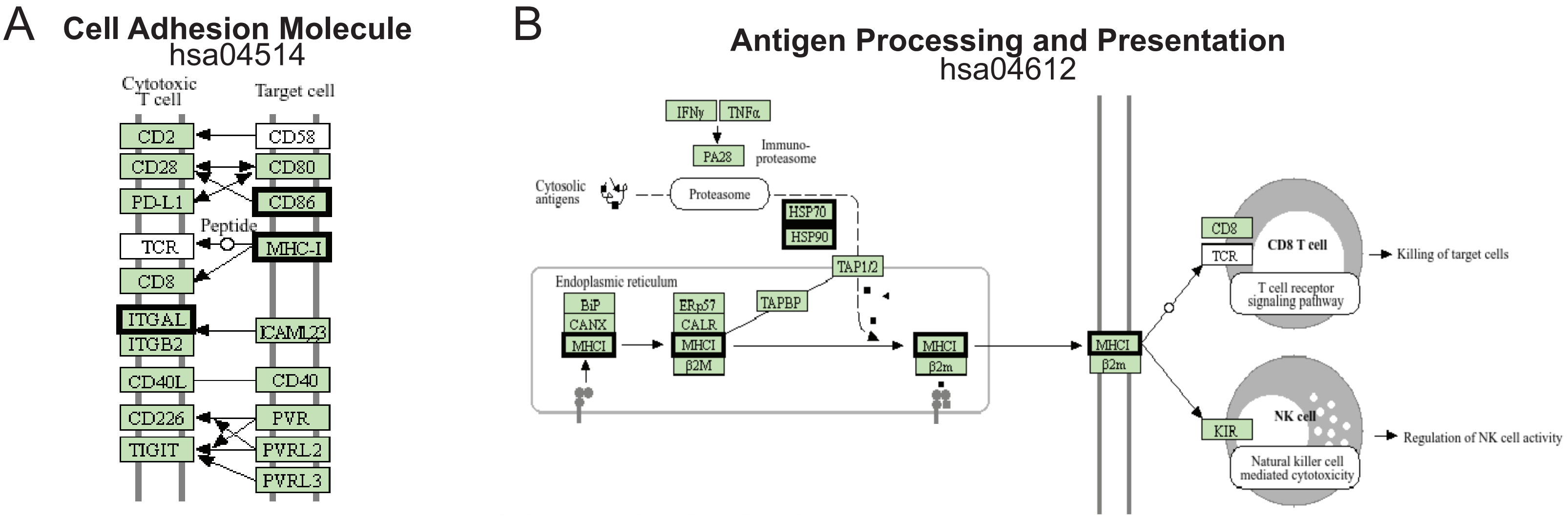
A
Cell Adhesion Molecule
hsa04514
Cytotoxic T cell
Target cell
CD2
CD58
CD28
CD80
PD-L1
CD86
Peptide
TCR
MHC-I
CD8
ITGAL
ITGB2
ICAM1,2,3
CD40L
CD40
CD226
PVR
TIGIT
PVRL2
PVRL3
B
Antigen Processing and Presentation
hsa04612
IFNγ
TNFα
PA28
Immuno-proteasome
Cytosolic antigens
Proteasome
HSP70
HSP90
TAP1/2
Endoplasmic reticulum
TAPBP
BiP
CANX
MHCI
ERp57
CALR
MHCI
β2M
MHCI
β2m
MHCI
β2m
CD8
TCR
CD8 T cell
T cell receptor signaling pathway
Killing of target cells
NK cell
KIR
Natural killer cell mediated cytotoxicity
Regulation of NK cell activity

**Supplementary Figure S12. (A-B)** Annotated KEGG pathway map highlighting the genes downregulated by CR alpha cells and within Cell Adhesion Molecule (A) and Antigen Processing and Presentation (B) pathways, respectively. Black bounding boxes mark CR downregulated genes in alpha cells compared to AL fed mice.

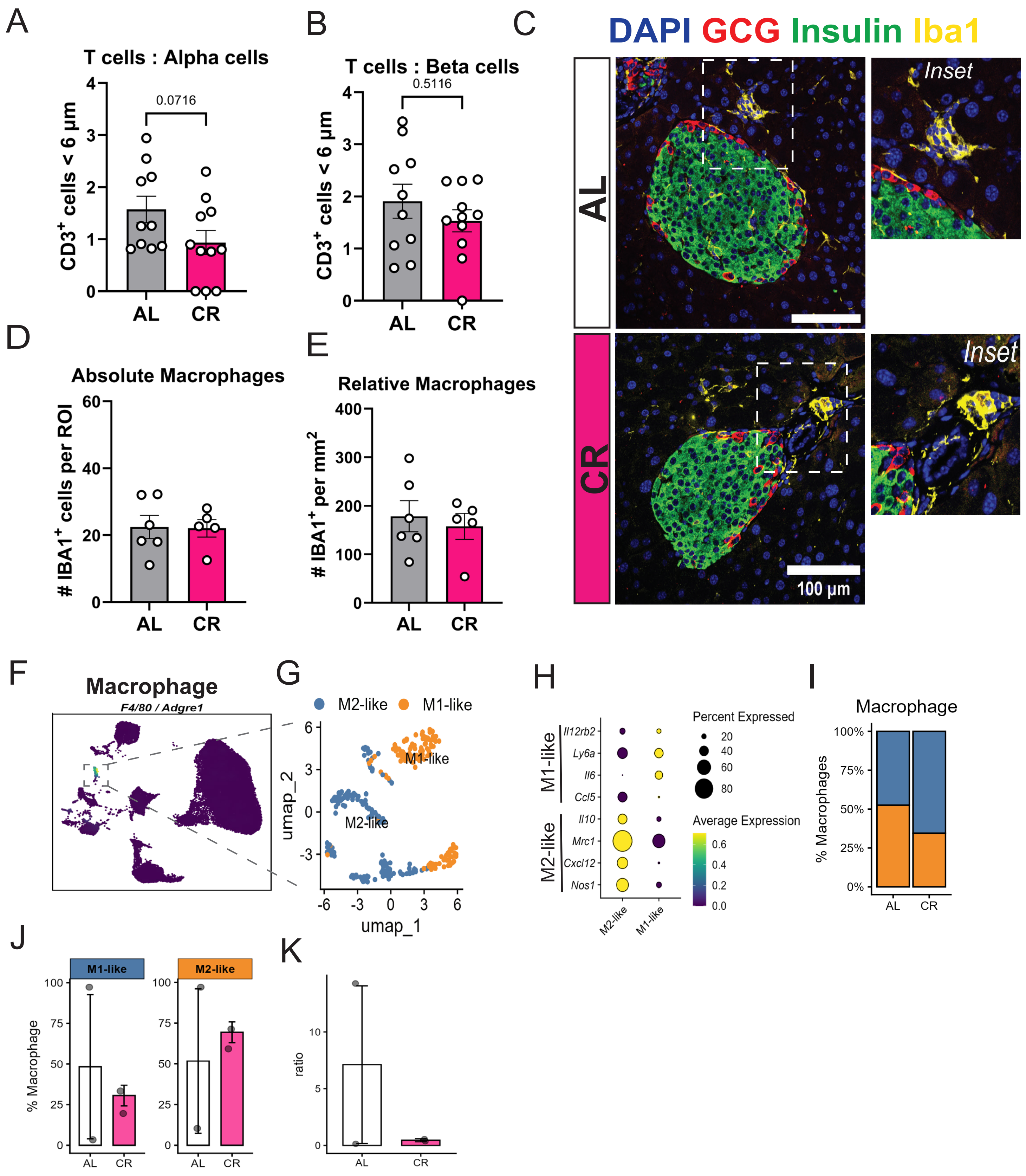
A
T cells : Alpha cells
0.0716
CD3+ cells < 6 µm
AL
CR
B
T cells : Beta cells
0.5116
C
DAPI GCG Insulin Iba1
Inset
100 µm
D
Absolute Macrophages
### IBA1+ cells per ROI
E
Relative Macrophages
### IBA1+ per mm2
F
Macrophage
F4/80 / Adgre1
G
M2-like
M1-like
umap_1
umap_2
H
Il12rb2
Ly6a
Il6
Ccl5
Il10
Mrc1
Cxcl12
Nos1
Percent Expressed
Average Expression
I
Macrophage
% Macrophages
J
% Macrophage
K
ratio

**Supplementary Figure S13. (A-B)** Number of CD3+ T cells within 6 microns or less from alpha cells or beta cells, respectively. **(C)** Representative confocal microscopy image of AL and CR mouse islets immunostained with anti-glucagon (GCG), anti-insulin, or the macrophage marker Iba1. **(D-E)** Number of Iba1+ macrophages within the imaging region of interest (islet + peri-islet space) and macrophage density, respectively. **(F-G)** Annotated UMAP of islet macrophage states identified in our dataset. **(H)** Macrophage state gene expression for M1 and M2-like macrophages. **(I-K)** Relative composition of the macrophage population in AL and CR islets.

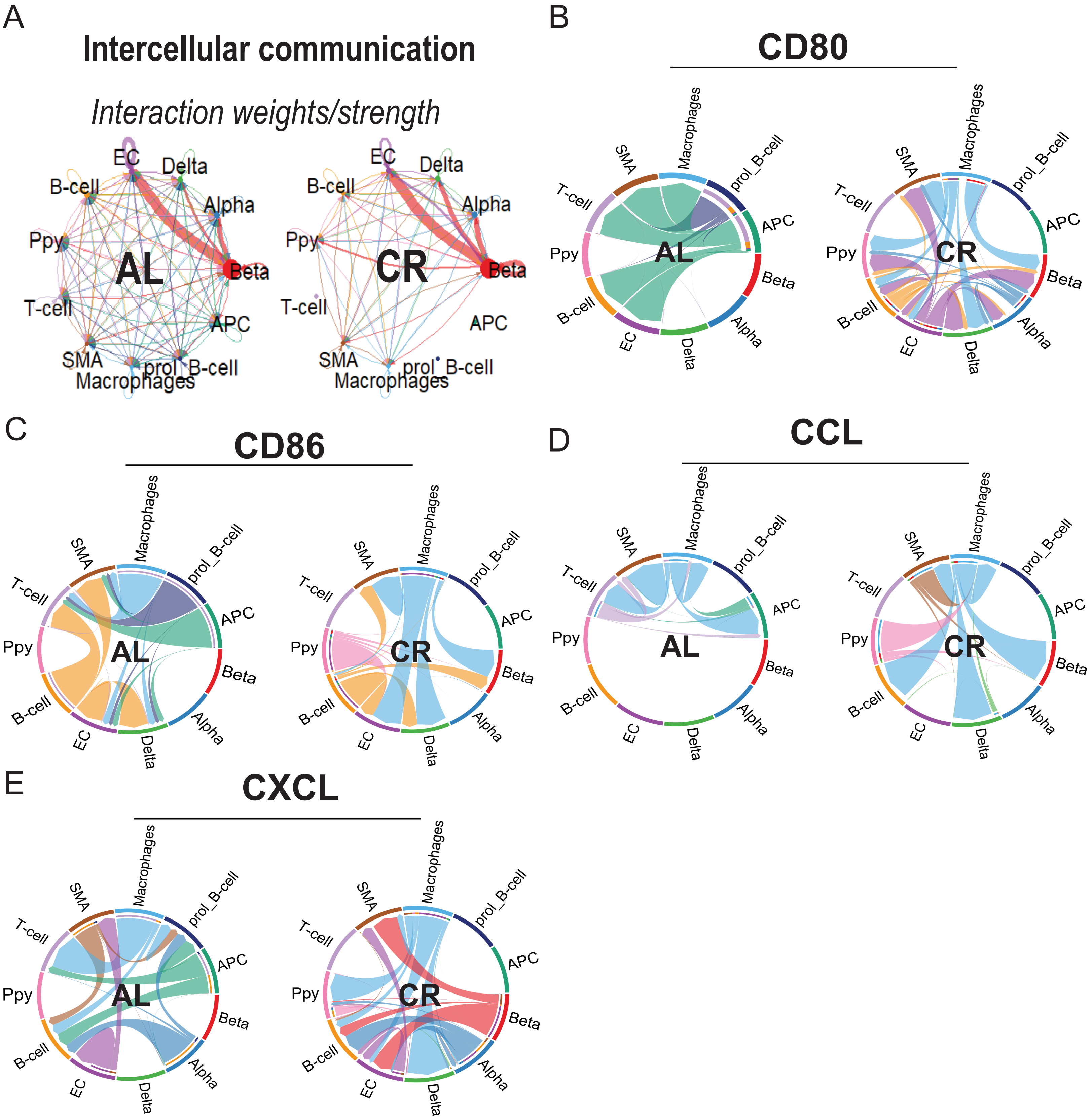
A
Intercellular communication
Interaction weights/strength
AL
CR
EC
Delta
B-cell
Alpha
Ppy
Beta
T-cell
APC
SMA
prol_B-cell
Macrophages
B
CD80
C
CD86
D
CCL
E
CXCL

**Supplementary Figure S14. (A)** Chord network plots representing the relative “interaction strength” index between identified cell types in AL and CR islets. **(B-E)** Chord network plots illustrating the cell-cell communication landscape for CD80, CD86, CCL, and CXCL signaling pathways in AL and CR islets.
